# Whole-brain *in vivo* base editing reverses autistic-like behaviors in mice

**DOI:** 10.1101/2022.01.25.477781

**Authors:** Weike Li, Jinlong Chen, Wanling Peng, Bo Yuan, Wenjian Han, Yiting Yuan, Zhenyu Xue, Jincheng Wang, Zhifang Chen, Shifang Shan, Shujia Zhu, Min Xu, Tianlin Cheng, Zilong Qiu

## Abstract

Autism spectrum disorder (ASD) is a highly heritable neurodevelopmental disorder associated with deficits in social communication and stereotypical behaviors. Numerous ASD-related genetic mutations have been identified and genome editing methods have been developed but successful genome editing in the whole-brain scale to alleviate autistic-like behaviors in animal models has not been achieved. Here we report the development of a new CRISPR-mediated cytidine base editor (CBE) system, which converts C·G base pairs to T·A. We demonstrate the effectiveness of this system by targeting an ASD-associated *de novo* mutation in the *MEF2C* gene (c.104T>C, p.L35P). We constructed a *Mef2c* L35P knock-in mouse and observed that *Mef2c* L35P heterozygous mice displayed autistic-like behaviors, including deficits in social behaviors and repetitive behaviors. We programmed the CBE to edit the C·G base pairs of the mutated *Mef2c* gene (c.104T>C, p.L35P) to T·A base pairs and delivered it via a single dose intravenous injection of blood brain barrier (BBB)-crossing AAV-PHP.eB vector into the mouse brain. This treatment restored MEF2C protein levels and reversed impairments in social interactions and repetitive behaviors in *Mef2c* L35P heterozygous mice. Together, this work presents an *in vivo* gene editing strategy in which correcting a single nucleotide mutation in the whole-brain scale could be successfully achieved, further providing a new therapeutic framework for neurodevelopmental disorders.

## Main text

Autism Spectrum Disorder (ASD) is a highly heritable neurodevelopmental disorder, characterized by deficits in social interaction and stereotypic behaviors ^1–3^. Rare *de novo* variants, including single nucleotide variants (SNVs) and copy number variants (CNVs), have been confirmed as important contributors to the pathogenesis of ASD ^4,5^.

Myocyte-specific enhancer factor 2C (MEF2C), a member of the MEF2 transcriptional factor family, was reported to be implicated in ASD, as recurrent *de novo* variants of the *MEF2C* gene were found in people diagnosed with ASD ^6,7^. MEF2C is abundantly expressed in the cortex, hippocampus and amygdala of adult mice ^8^ and plays a vital role in neuronal differentiation, neural development and synaptic plasticity ^9–14^. Microdeletions of the *MEF2C*-containing chromosomal segment (5q14.3-q15) causes developmental deficits in children, including intellectual disability (ID), poor reciprocal behaviors, lack of speech, stereotypic and repetitive behavior, and epilepsy ^15–17^, suggesting that *MEF2C* haploinsufficiency causes severe defects of brain development.

The Clustered regularly interspaced short palindromic repeats (CRISPR) and CRISPR-associated protein 9 (Cas9) system has been widely used for genome editing ^18,19^. With well-designed single guide RNA (sgRNA), double-stranded DNA breaks (DSBs) are introduced into genomic targets via Cas9-mediated cleavage, followed by either non-homologous end-joining (NHEJ) or homology-directed repair (HDR) pathway ^20^. Although several approaches increasing HDR efficiencies have been developed to repair diseases-causing genetic mutations ^20^, induced DSBs may lead to unexpected genomic instability. Meanwhile, base editors (BEs), which edit single base pairs precisely without generation of DSBs, have been developed through fusion of Cas9 nickase with various deaminases ^21–23^. Cytidine and adenosine deaminases have been used to generate cytidine base editors (CBEs) for C-T conversion and adenosine base editors (ABEs) for A-G conversion, respectively ^21,22^. Take advantage of adeno-associated virus (AAV)-mediated delivery, *in vivo* base editing has been applied to various disease models ^24,25^. Nevertheless, whether base editors can be applied for neurodevelopmental disorders remains to be addressed.

In this work, we identified a *de novo* SNV in the *MEF2C* gene (c.104T>C, p.L35P) from people with ASD. We then constructed *Mef2c* L35P knock-in mouse and observed that *Mef2c* L35P heterozygous mice displayed autistic-like behaviors, such as social deficits and repetitive behavior. MEF2C protein expression in the brain of *Mef2c* L35P heterozygous mice was markedly reduced compared to WT mice, suggesting that the L35P mutation may decrease structural stability or accelerate MEF2C protein degradation.

We combined the newly developed CBE with the blood brain barrier (BBB)-crossing AAV-PHP.eB system and delivered CBE into the mouse brain by intravenous injection ^26,27^. This single CBE-mediated base editing treatment was sufficient to restore MEF2C protein levels and correct the defects in social interactions and repetitive behaviors of *Mef2c* L35P heterozygous mice.

### Identification of a *de novo* SNV in the *MEF2C* gene in a Chinese patient with ASD

We performed whole-exome sequencing of one ASD patient with unaffected parents collected in the Xinhua hospital affiliated to Shanghai Jiao Tong University School of Medicine, and identified a *de novo* SNV in the *MEF2C* gene (c.104T>C, p.L35P) (Fig. 1a and Fig. S1a), which we validated by Sanger sequencing (Fig. 1b). The amino acid change (L35P) caused by the *de novo* variant is located in the MCM1, AGAMOUS, DEFICIENS, and SRF (MADS) domain of MEF2C protein (Fig. 1c, S1b). This variant (*MEF2C*, chr5: 88804752) is not present in over 18,800 genomes from East Asian populations in the gnomAD database (http://gnomad.broadinstitute.org), indicating that it is a rare variant. In a recent report including 112 Chinese patients with intellectual disability and Rett-like symptoms, researchers identified numerous *de novo* and inherited variants in *MEF2C*, suggesting that *MEF2C* is a critical risk gene for developmental disorders in the Chinese population^28^. A schematic illustration of various locations of genetic mutations in the MEF2C protein is shown in Fig. 1c ^29–33^.

**Figure 1.**
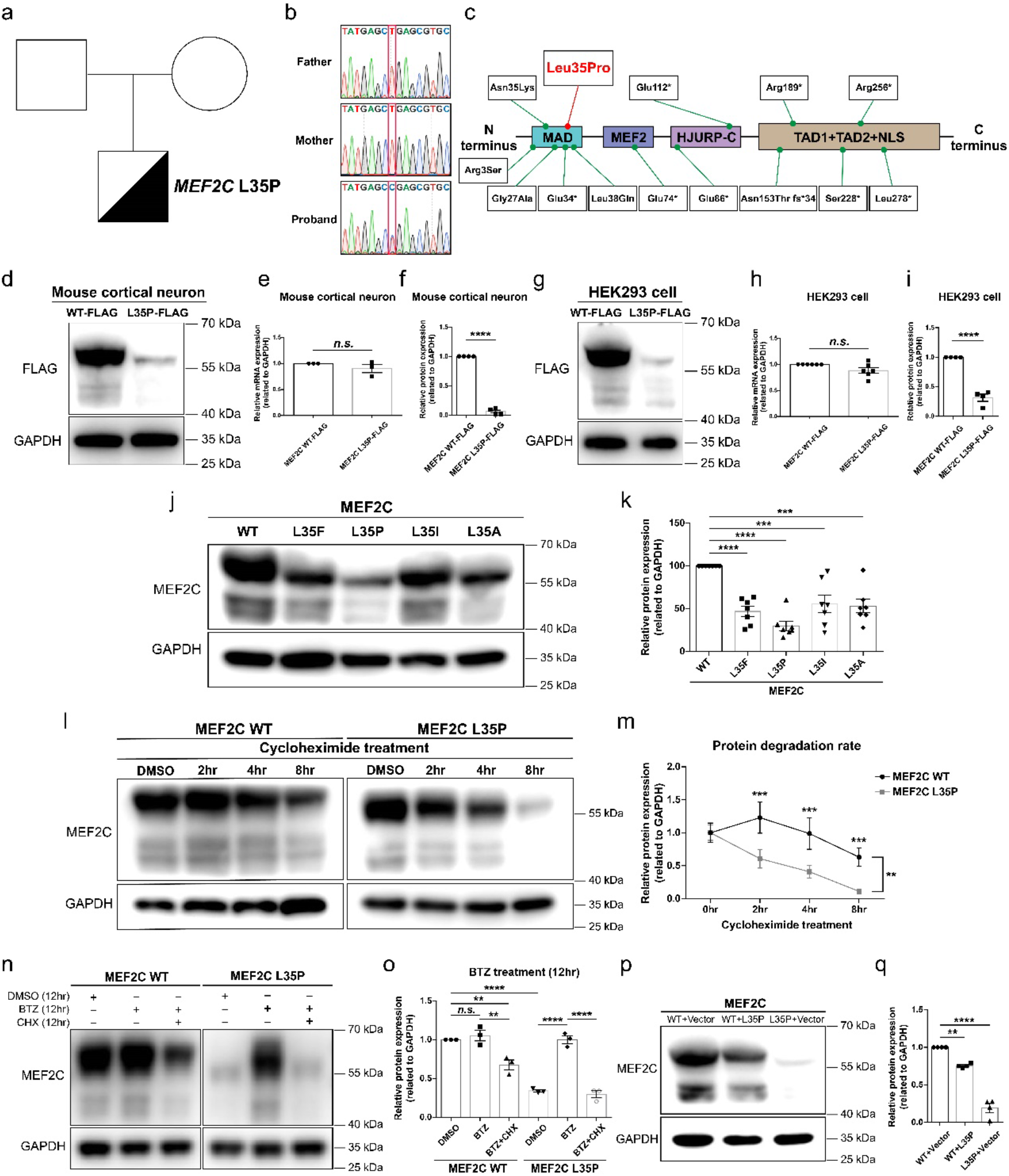
A *de novo* mutation in the *MEF2C* gene associated with ASD causes rapid degradation of MEF2C protein. **a**, Schematic illustration of the *de novo MEF2C* variant identified in a Chinese ASD simplex family (Squares and circles represent males and females; blank represent non-carrier; Half blank and half black represent carrying the *de novo* variant in the *MEF2C* gene). **b**, Validation of the *de novo* variant of *MEF2C* by sanger sequencing (Red box indicates the variant). **c**, The location of the *de novo* variant (L35P) in the MEF2C protein. Green dots indicate 13 SNVs on *MEF2C* exons reported previously; MADS: MCM1, Agamous, Deficiens, Serum response factor; MEF2, myocyte enhancer factor 2; TAD, transcriptional activation domain; NLS, nuclear location signal. **d**, Western blotting of Flag-tagged MEF2C WT and L35P expressed in cultured primary cortical neurons. Quantitative analysis of mRNA level (**e**) and protein level (**f**) of Flag-tagged MEF2C WT and L35P expressed cultured primary cortical neurons normalized to GAPDH (*n* = 3). **g**, Western blotting of Flag-tagged MEF2C WT and L35P expressed in HEK293 cells (*n* = 6). Quantitative analysis of mRNA level (**h**) and protein level (**i**) of Flag-tagged MEF2C WT and L35P expressed in HEK293 cells. (unpaired two-tailed Student’s *t*-test). **j**, Western blotting of MEF2C WT, L35F, L35P, L35I or L35A expressed in transfected HEK293 cells. **k**, Quantification of relative MEF2C protein levels normalized to GAPDH (*n* = 7, one-way ANOVA). **l**, Western blotting of *MEF2C* WT and L35P expressed in HEK293 cells, with cycloheximide (CHX) treatment for indicated time before immunoblotting analysis. **m**, Quantitative analysis of relative protein expression of *MEF2C* WT and L35P in (l). (*n* = 4, two-way ANOVA). **n**, Western blotting of MEF2C WT and L35P expressed in HEK293 cells, with bortezomib (BTZ) or CHX treatment for 12 hr before immunoblotting analysis. **o**, Quantification of relative MEF2C protein levels normalized to GAPDH in (**n**) (*n* = 3, one-way ANOVA). **p**, Western blotting of MEF2C-WT or MEF2C-L35P co-transfected with either empty vector or L35P in HEK293 cells. **q**, Quantitative analysis of Western blotting in (**p**) (*n* = 4, one-way ANOVA). Each dot in western blotting represents one independent experiment. *n* is biological repeat numbers of independent experiments. Statistical values represent the mean ± s.e.m. *\*\* p* < 0.01, *** *p* <0.001*\*\*\*\* p* < 0.0001.

To investigate whether the L35P mutation may affect the proper function of the MEF2C protein, we examined the expression level of flag-tagged wild-type (WT) and L35P MEF2C protein in both mouse cortical neurons and HEK293 cells. Intriguingly, we found that the protein level of MEF2C-L35P was significantly lower compared to MEF2C-WT (Fig. 1d-1i), whereas mRNA levels of WT and L35P MEF2C remained the same (Fig. 1e, 1h), suggesting that the L35P mutation may affect the level of MEF2C protein.

To further investigate the importance of L35 for MEF2C protein stability, we tested the impact of other hydrophobic amino acids such as isoleucine or phenylalanine (I/F), as well as alanine (A). L35F, L35I and L35A all led to decreased levels of MEF2C proteins, indicating that L35 is critical for MEF2C protein stability (Fig. 1j, 1k). After cycloheximide (CHX) treatment, an inhibitor of protein translation ^34^, levels of MEF2C-L35P decreased faster than MEF2C-WT, indicating that the L35P mutation accelerated MEF2C protein degradation (Fig. 1l, m). Treatment with the proteasome inhibitor bortezomib (BTZ) restored levels of MEF2C-L35P, suggesting that the rapid degradation of MEF2C-L35P was mediated by the ubiquitin-dependent pathway (Fig. 1o). Finally, to investigate whether the MEF2C-L35P protein may affect the protein level of MEF2C-WT, we co-transfected MEF2C-WT along with MEF2C-L35P into HEK293 cells. Co-expression of MEF2C-L35P with MEF2C-WT led to marked reduction of total MEF2C protein level (Fig. 1p, 1q), suggesting that the L35P mutation exhibits a dominant negative effect on MEF2C.

### MEF2C-L35P leads to aberrant neuronal dendritic and axonal development

To investigate the impact of MEF2C-L35P on neurons, we constructed a short hairpin RNA (shRNA) specifically targeting the mouse *Mef2c* gene. With two designed shRNA candidates (Fig. S2a), we assessed the knock-down efficiency by examining endogenous *Mef2c* mRNA levels in mouse cortical neurons. sh *Mef2c*-1 exhibited higher efficiency and was used in subsequent studies (Fig. S2b). We found that expression of endogenous Mef2c protein was effectively reduced by sh *Mef2c*-1 (Fig. S2c, d).

Consistent with a previous report ^35^, we found that knockdown of *Mef2c* led to decreased dendritic length and branch numbers, as well as axon length (Fig. S3a-f), which could be fully restored by co-transfection of a shRNA-resistant MEF2C-WT construct, but not MEF2C-L35P (Fig. S3a-f). These results suggest that the L35P mutation impairs MEF2C function in neurons.

### Abnormal neural development and autistic-like behaviors in *Mef2c* L35P knock-in mice

To investigate the role of MEF2C-L35P in ASD pathogenesis, we constructed *Mef2c* L35P knock-in mice by CRISPR/Cas9-mediated gene targeting (Fig. S4a). MEF2C protein levels of *Mef2c* L35P^+/−^ mice were markedly decreased compared to WT mice (Fig. 2a-c). Since MEF2C is widely expressed in the cortex, hippocampus and amygdala ^8,36^, we performed immunohistochemical staining to examine the expression of MEF2C in the brain of *Mef2c* L35P^+/−^ mice. We found that the fluorescence intensity of MEF2C signals in *Mef2c* L35P^+/−^ outer cortex, dentate gyrus and amygdala were reduced compared to WT mice (Fig. 2d, Fig. S4b-d).

**Figure 2.**
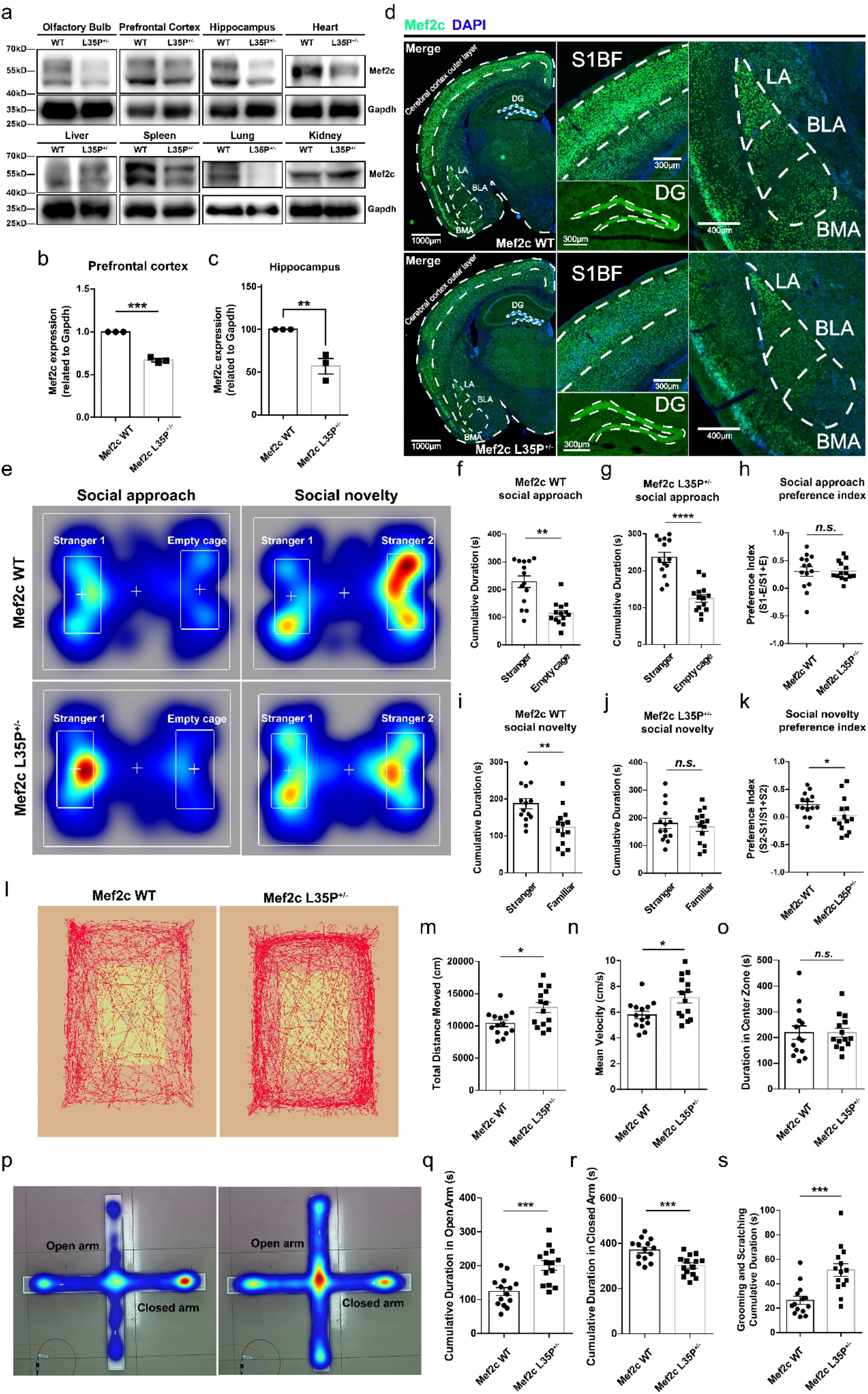
Reduced MEF2C protein levels and autistic-like behaviors in *Mef2c* L35P knock-in mice. **a**, Western blotting of MEF2C protein in various brain regions and peripheral tissues extracted from *Mef2c* WT and L35P^+/−^ mice. Quantitative analysis of relative Mef2c expression of PFC (**b**) and Hippocampus (**c**) in Mef2c WT and L35P^+/−^ mice (*n* = 3). **d**, Immunohistochemical staining for Mef2c (green) and DAPI (nuclei marker, blue) in the cerebral cortex, hippocampus and amygdala of *Mef2c* WT and L35P^+/−^ mice (S1BF, somatosensory cortex 1 barrel field; DG, dentate gyrus; LA, lateral amygdala; BLA, basolateral amygdala; BMA, basomedial amygdala; white dotted box indicates corresponding brain region). Scale bars, 1000μm (left); 300μm (middle); 400μm (right). **e**, Representative heat maps for locomotion of *Mef2c* WT and L35P^+/−^ mice on the social approach and social novelty session in the three-chamber test. Quantification of cumulative duration of interacting with mouse and empty cage of *Mef2c* WT (**f**) and L35P^+/−^ mice (**g**) (*n* = 14 for each group). **h**,The preference index of social approach test for *Mef2c* WT and L35P^+/−^ mice. Quantification of cumulative duration of interacting with novel stranger mouse and familiar mouse of *Mef2c* WT (**i**) and L35P^+/−^ (**j**) mice (*n* = 14 for each group). **k**,The preference index of social novelty test for *Mef2c* WT and L35P^+/−^ mice. **l**, Representative locomotion track traces of *Mef2c* WT and L35P^+/−^ mice in the open field test. Quantitative analysis of total distance (**m**), mean velocity (**n**), and duration in the center zone (**o**) of *Mef2c* WT and L35P^+/−^ mice in the open field test (*n* = 14 for each group). **p**, Representative heat maps of *Mef2c* WT and L35P^+/−^ mice in the elevated plus maze (open arm indicates top and bottom arms; closed arm indicates left and right arms). Quantification of cumulative duration exploring in the open arm (**q**) and closed arm (**r**) of *Mef2c* WT and L35P^+/−^ mice (*n* = 14 for each group). **s**, Quantitative analysis of cumulative self-grooming and scratching time of *Mef2c* WT and L35P^+/−^ mice (*n* = 14 for each group). *n* represents mice number. Statistical values represent the mean ± s.e.m. *\*p* < 0.05, *\*\*p* < 0.01, *\*\*\*p* < 0.001, *\*\*\*\*p* < 0.0001, unpaired two-tailed Student’s *t*-test.

Multiple ASD animal models exhibit aberrant inhibitory interneuron development ^37^, and it was reported that the population of parvalbumin (PV) positive interneurons decreased in the hippocampus of *Mef2c* ^+/−^ mice ^35,38,39^. Thus, we examined PV-positive GABAergic neurons in the brain of *Mef2c* L35P^+/−^ mice. By immunohistochemical staining of parvalbumin, we found that there was a prominent reduction of PV-positive interneurons in the retrosplenial cortex (RSC), dentate gyrus (DG), somatosensory cortex (SC) and visual cortex (VC) of *Mef2c* L35P^+/−^ mice compared to WT mice (Fig. S4e-i). In contrast, *Mef2c* L35P^+/−^ mice showed normal populations of somatostatin positive interneurons in RSC, Hip and SC, suggesting that MEF2C dysfunction specifically impaired development of PV-positive interneurons (Fig. S4j-m). The aberrant development of PV-positive GABAergic neurons suggests that an imbalance of excitatory/inhibitory (E/I) synaptic transmission may exist in the brain of *Mef2c* L35P^+/−^ mice, which may contribute to ASD pathogenesis ^40,41^.

### *Mef2c* L35P^+/−^ mice display autistic-like and *Mef2c* haploinsufficiency syndrome (MCHS)-like behaviors

We next examined whether MEF2C-L35P mutation affected the gross development of the mice by measuring body weights of WT and *Mef2c* L35P^+/−^ mice from birth to 9 weeks old. We found there was no difference in body weights between WT and *Mef2c* L35P^+/−^ mice (Fig. S5a). Previous reports have shown that *Mef2c*^+/−^ mice exhibited various abnormal behaviors, including deficits in social interaction, repetitive behaviors and hyperactivity ^13,35^. Interestingly, using the classic three-chamber test, we found that *Mef2c* L35P^+/−^ mice exhibited normal social approach but abnormal performance in the social novelty test (Fig. 2e-k) compared to WT mice (Fig. 2e, f, i). In the novel object recognition test, *Mef2c* L35P^+/−^ mice showed similar preference for novel object over familiar object compared to WT mice, suggesting that *Mef2c* L35P^+/−^ mice have a specific defect in recognizing novel partners (Fig. S5b-d).

*Mef2c* L35P^+/−^ mice did not display anxiety-like phenotypes, but exhibited remarkable hyperactivity in the open field test (Fig. 2l-o). In the elevated plus maze test, we found that *Mef2c* L35P^+/−^ mice exhibited more preference for open arms rather than closed arms, suggesting that *Mef2c* L35P^+/−^ mice showed hyperactivity rather anxiety-like phenotype (Fig. 2p-r). *Mef2c* L35P^+/−^ mice also exhibited prominent repetitive behavior, showing significantly more self-grooming and scratching than WT mice (Fig. 2s).

Lastly, we examined whether *Mef2c* L35P^+/−^ mice have normal spatial learning and memory capability with Barnes maze. We found that *Mef2c* L35P^+/−^ mice exhibited the same learning curve during training session and cumulative duration within the target zone in the test session compared to WT mice, indicating that *Mef2c* L35P^+/−^ mice have normal ability for learning and memory for spatial information (Fig. S5e-g).

### Establishment of the new CBE for correcting the MEF2C-L35P mutation

To correct the *Mef2c* L35P mutation (c.104T>C), CBEs are required to convert mutated C·G base pairs to T·A. Based on the existing CBEs ^21,22^, we designed a new CBE tool derived from SpG, a *Streptococcus pyogenes* Cas9 variant targeting NGN protospacer-adjacent motif (PAM) ^42^, with human cytidine deaminase APOBEC3A-Y130F with minimal RNA off-targeting effects ^43,44^, and uracil glycosylase inhibitor (UGI) (Fig. 3a). To maximize base editing efficiency, we fused cytidine deaminase inside Cas9 ^26,45^. Then we designed two sgRNAs (sgRNA-C8 and sgRNA-C15) targeting the mutation site in *Mef2c* gene (Fig. 3b).

**Figure 3.**
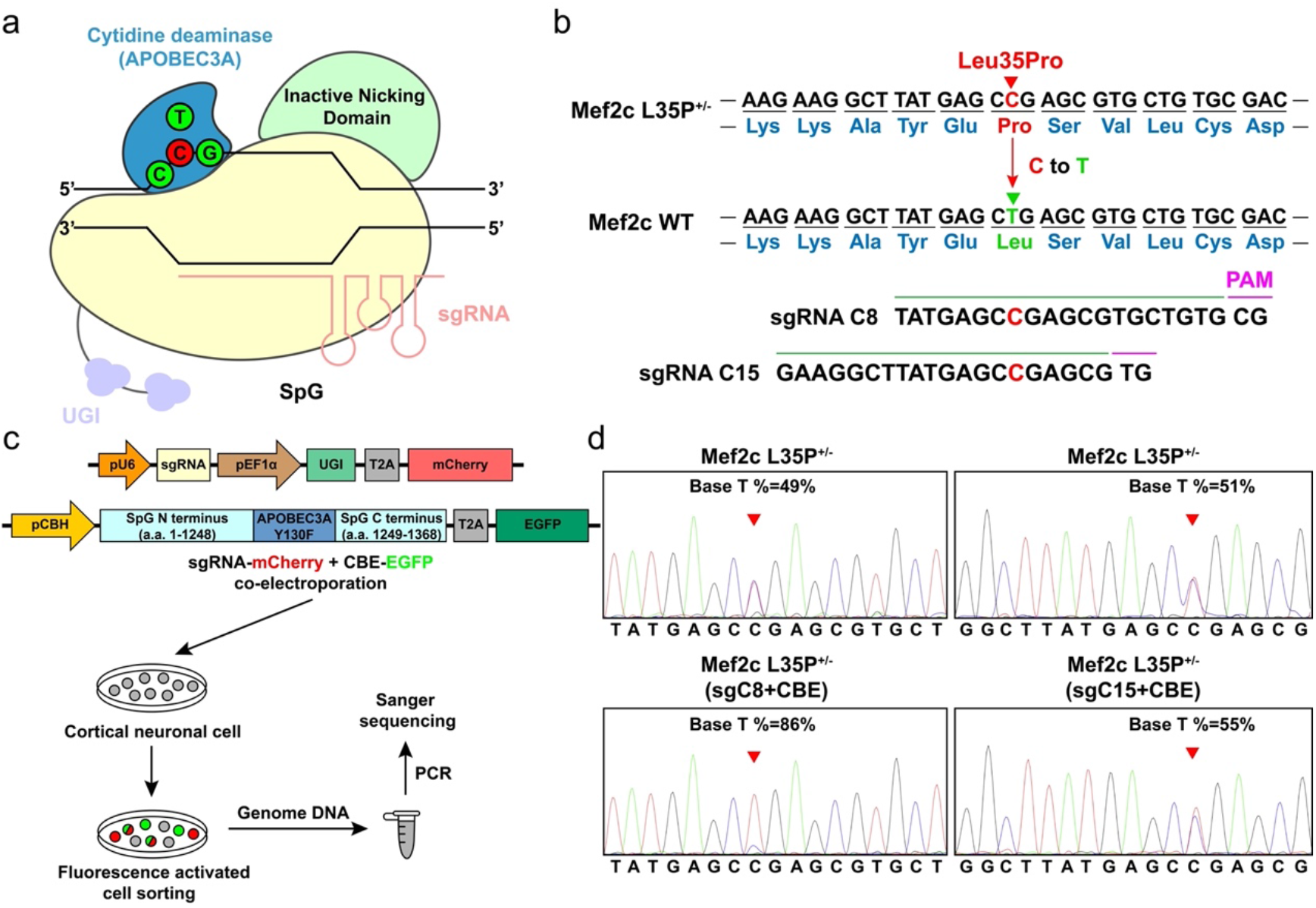
Design and validation of the CBE gene editing system. **a**, Schematic diagram of CRISPR/Cas9-based CBE gene editing system, including SpG, sgRNA, UGI and APOBEC3A-Y130F (red circle indicates the mutation site, green circles indicate correct base pair). **b**, Design of sgRNAs targeting to the mutation site in the mouse Mef2c gene. **c**, Schematic flow for validating the efficiency of the CBE system in vitro in mouse cortical neurons. Cultured mouse cortical neurons were co-electroporated with sgRNA-mCherry and CBE-EGFP plasmids followed by sorting with both mCherry and GFP signals. **d**, Representative sanger sequencing results of primary cortical neurons from Mef2c L35P+/− mice after CBE gene editing (red arrowhead indicate gene loci without gene editing (upper panels) and with SgC8 (lower left panel), SgC15 (lower right panel). The percentage of T (c.104T) is obtained by calculating T base ratio (T /(C+T) * 100%).

To evaluate the efficiency of the newly developed CBE system in post-mitotic neurons, we co-transfected CBE with two sgRNAs into cultured primary cortical neurons from *Mef2c* L35P^+/−^ mice. Neurons transfected with both sgRNA and CBE were collected with fluorescence activated cell sorting (FACS) to evaluate the mutation status (c.104T>C) by PCR and Sanger sequencing (Fig. 3c). We found that the wild-type T percentage (86%) edited with sgRNA-C8 (SgC8) and CBE is significantly higher than that in the negative control (49%), while the T percentage edited with sgRNA-C15 and CBE remained similar (55%) to negative control (51%) (Fig. 3d). Therefore, sgRNA-C8 was chosen for subsequent *in vivo* therapeutic base editing of *Mef2c* L35P^+/−^ mice.

### *In vivo* base editing mediated by AAV in *Mef2c* L35P^+/−^ mice

Because of the limited packaging capacity of AAV, we used an intein-mediated split strategy to generate dual-AAV system ^24,46^, with one AAV containing the N-terminus of SpG (a.a. 1-793) and sgRNA-C8 expression cassette, and the other AAV containing the C-terminus of SpG-APOBEC3A-UGI (a.a. 794-1368) (Fig. 4a). To deliver the CBE system into the mouse brain, we used the BBB-crossing adeno-associated virus (AAV-PHP.eB) and delivered the AAVs by tail veins injection in 1-month old *Mef2c* WT or L35P^+/−^ mice (Fig. 4a) ^24,27^.

**Figure 4.**
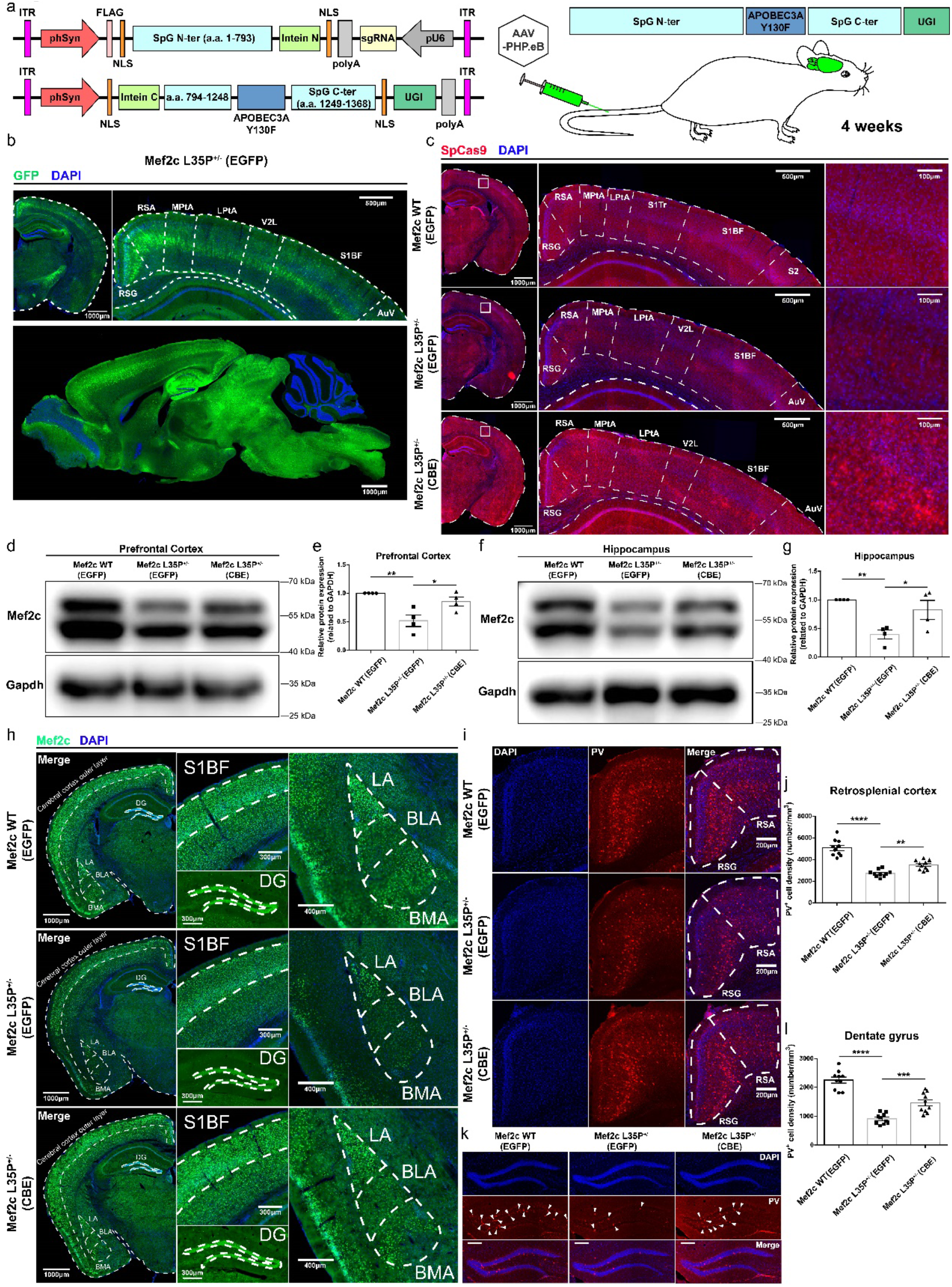
Restoration of Mef2c protein level and abnormal inhibitory interneuron development in *Mef2c* L35P^+/−^ mice after the CBE gene editing *in vivo*. **a**, Schematic illustration of dual AAV plasmids engineered by intein-dependent protein split strategy. **b**, Representative immunofluorescent staining for EGFP (green) and DAPI (blue) to determine the AAV infection efficiency in the brain through intravenous injection in the tail vein (RSG, retrosplenial granular cortex; RSA, retrosplenial agranular cortex; MPtA, medial parietal association cortex; LPtA, lateral parietal association cortex; V2L, secondary visual cortex, lateral area; AuV, secondary auditory cortex, ventral area). Scale bars, 1000μm (upper left and bottom); 500μm (upper right). **c**, Representative immunofluorescent staining for spCas9 (red) and DAPI (blue) to examine CBE delivery efficiency in the brain (S1Tr, primary somatosensory cortex, trunk region; S2, secondary somatosensory cortex; white dotted box indicates corresponding brain region). Scale bars, 1000μm (left); 500μm (middle); 100μm (zoom magnification, right). Representative western blotting of prefrontal cortex (**d**) and hippocampus (**f**) extracted from *Mef2c* WT mice treated with AAV-EGFP, and *Mef2c* L35P^+/−^ mice injected with either EGFP or CBE. **e**, **g**,Quantification of western blotting in (**d**) and (**e**), respectively (*n* = 4 mice per group). **h**, Immunohistochemical staining for Mef2c (green) and DAPI (blue) in the cerebral cortex outer layer, hippocampus and amygdala of *Mef2c* WT mice injection of AAV-EGFP, and *Mef2c* L35P^+/−^ mice injected with either EGFP or CBE. Scale bars, 1000μm (left); 300μm (middle); 400μm (right). **i**,**k**, Immunohistochemical staining for PV (red), Mef2c (green) and DAPI (blue) in the retrosplenial cortex and dentate gyrus of *Mef2c* WT mice injection of AAV-EGFP, and *Mef2c* L35P^+/−^ mice injected with either EGFP or CBE.. Scale bars, 200μm., Quantification of PV-positive cell density in the retrosplenial cortex (**j**) and dentate gyrus (**l**) (*n*=10 brain slices from 4 mice per group). Statistical values represent the mean ± s.e.m. *\*p* < 0.05, *\*\*p* < 0.01, *\*\*\*p* < 0.001, *\*\*\*\*p* < 0.0001, unpaired two-tailed Student’s *t*-test.

We evaluated the delivery efficiency of AAV-PHP.eB vectors using AAV-hSyn-EGFP, and immunohistochemical analysis 6-8 weeks after intravenous injection. We found that the expression of EGFP was widely distributed in the mouse brain, especially in cortical, hippocampal and midbrain regions (Fig. 4b). To assess the expression efficiency of the dual-AAV CBE system in the brain of *Mef2c* L35P^+/−^ mice, we performed immunohistochemical staining with an antibody against SpCas9, which could recognize the SpCas9 variant SpG, and found that SpG was expressed in the cortical regions as well as hippocampus (Fig. 4c).

To examine whether the CBE system may lead to base editing *in vivo*, we collected the hippocampal tissue from mice injected with either AAV-PHP.eB-EGFP or AAV-PHP.eB-CBE and amplified the target segments containing *Mef2c* (c.104T>C). After performing Sanger sequencing, we found that the percentage of targeted T in the CBE injection group was around 55%, whereas it in the EGFP injection group is 51%. Although the change is not dramatic, we reasoned that around 15% of cells in the mouse hippocampus are neurons, and MEF2C is highly expressed in neurons but not non-neuronal cells. Since base editing mediated by the new CBE system should preferentially function in the neurons due to driven by the human synapsin 1 promoter (*hSyn*), it may be difficult to observe large changes in percentage of targeted T collected from mixed brain tissues. We further evaluated the off-targeting effects by examining the C-to-T editing frequency in the 11 potential off-target sites (OT1-OT11) and observed minor changes in one of off-targeting sites (OT4) (Fig. S6b), suggesting that off-targeting effects in the new CBE system are minimal.

In order to validate the editing efficiency of this new CBE system, we further injected the dual-AAV CBE system with AAV9 vector into one side of hippocampus of *Mef2c* L35P^+/−^ mice, in which the contralateral side of hippocampus injected with AAV-EGFP as a negative control (Fig. S6c). After extraction of genomic DNA from hippocampus 4 weeks after injection, we amplified the target segments with PCR. Sanger sequencing results revealed that the T percentage from the dual-AAV CBE injection side (54%) was higher than the other side (50%), indicating that dual-AAV CBE system successfully achieved C-T conversion in post-mitotic neurons (Fig. S6d).

We next investigate whether the decreased MEF2C protein level in *Mef2c* L35P^+/−^ mice could be rescued after *in vivo* base editing by CBE, we performed immunoblot with brain lysates collected from WT or *Mef2c* L35P^+/−^ mice injected with either AAV-EGFP or AAV-CBE. We found that dual-AAV CBE successfully restored MEF2C protein in prefrontal cortex and hippocampus of *Mef2c* L35P^+/−^ mice to levels comparable to WT mice (Fig. 4d-g). Immunohistochemistry also demonstrated that the endogenous MEF2C protein level was significantly rescued in cortical regions and amygdala of *Mef2c* L35P^+/−^ mice (Fig. 4h, Fig. S7a-c).

Furthermore, PV-positive neurons in various brain regions, such as retrosplenial cortex and dentate gyrus of *Mef2c* L35P^+/−^ mice were also restored after *in vivo* base editing with dual-AAV CBE, indicating that the imbalance of excitatory/inhibitory synaptic transmission in mutant mice may be recovered (Fig. 4i-l)^47^, even in adult mice. Intriguingly, the decreased level of PV-positive neurons in the visual cortex and somatosensory cortex were not restored after AAV-CBE injection, suggesting that the restoration of excitatory/inhibitory synaptic functions in the brain of *Mef2c* L35P^+/−^ mice might be region-specific (Fig. S7d-f).

To assess whether injection of AAV-CBE could rescue synaptic function in *Mef2c* L35P^+/−^ mice, we used a sparse labeling method to label neurons in medial prefrontal cortex (mPFC) of WT and *Mef2c* L35P^+/−^ mice with injection of GFP or CBE (Fig. S8a, b). We found that the density of mushroom spines and total spines in the mPFC of *Mef2c* L35P^+/−^ mice were markedly decreased compared to WT mice, which was significantly rescued by CBE, suggesting that impaired excitatory synaptic functions are largely rescued in the *Mef2c* L35P^+/−^ mice with the dual-AAV CBE system (Fig. S8c-e)

### *In vivo* base editing corrects autistic-like behaviors of *Mef2c* L35P^+/−^ mice

Finally, to test if autistic-like behaviors of *Mef2c* L35P^+/−^ mice could be rescued by *in vivo* base editing with the dual-AAV CBE system, we performed behavioral analysis. We conducted the three-chamber test for WT or *Mef2c* L35P^+/−^ mice four weeks after AAV injection. In general, AAV injection did not affect the social approach for WT or *Mef2c* L35P^+/−^ mice (Fig. 5a-e), however, the impaired social novelty observed in *Mef2c* L35P^+/−^ mice was fully rescued as compared to WT mice (Fig. 5f-i). We conducted another paradigm of social interaction behaviors, the social intruder test, for WT or *Mef2c* L35P^+/−^ mice treated with either AAV-CBE or AAV-GFP. This test measures reciprocal interaction time of mice actively interacting with a strange partner for 4 consecutive trials, followed by interacting with a new strange partner for the 5^th^ trial (Fig. 5j, 5k). We found that the decreased initial social interaction time during the first trial between *Mef2c* L35P^+/−^ mice and partners was fully rescued by the dual-AAV CBE system, further demonstrating that behavioral impairments can be rescued by *in vivo* CBE postnatally (Fig. 5j). On the fifth trial, we found that *Mef2c* L35P^+/−^ mice injected with AAV-EGFP had severe defects in recognizing familiar or stranger partners, compared to WT mice injected with AAV-EGFP (Fig. 5k-o). However, the abnormal social interaction of *Mef2c* L35P^+/−^ mice was fully rescued after delivery of *in vivo* CBE (Fig. 5k-o), indicating that correction of the *Mef2c* mutations *in vivo* can restore the normal social behaviors in mice.

**Figure 5.**
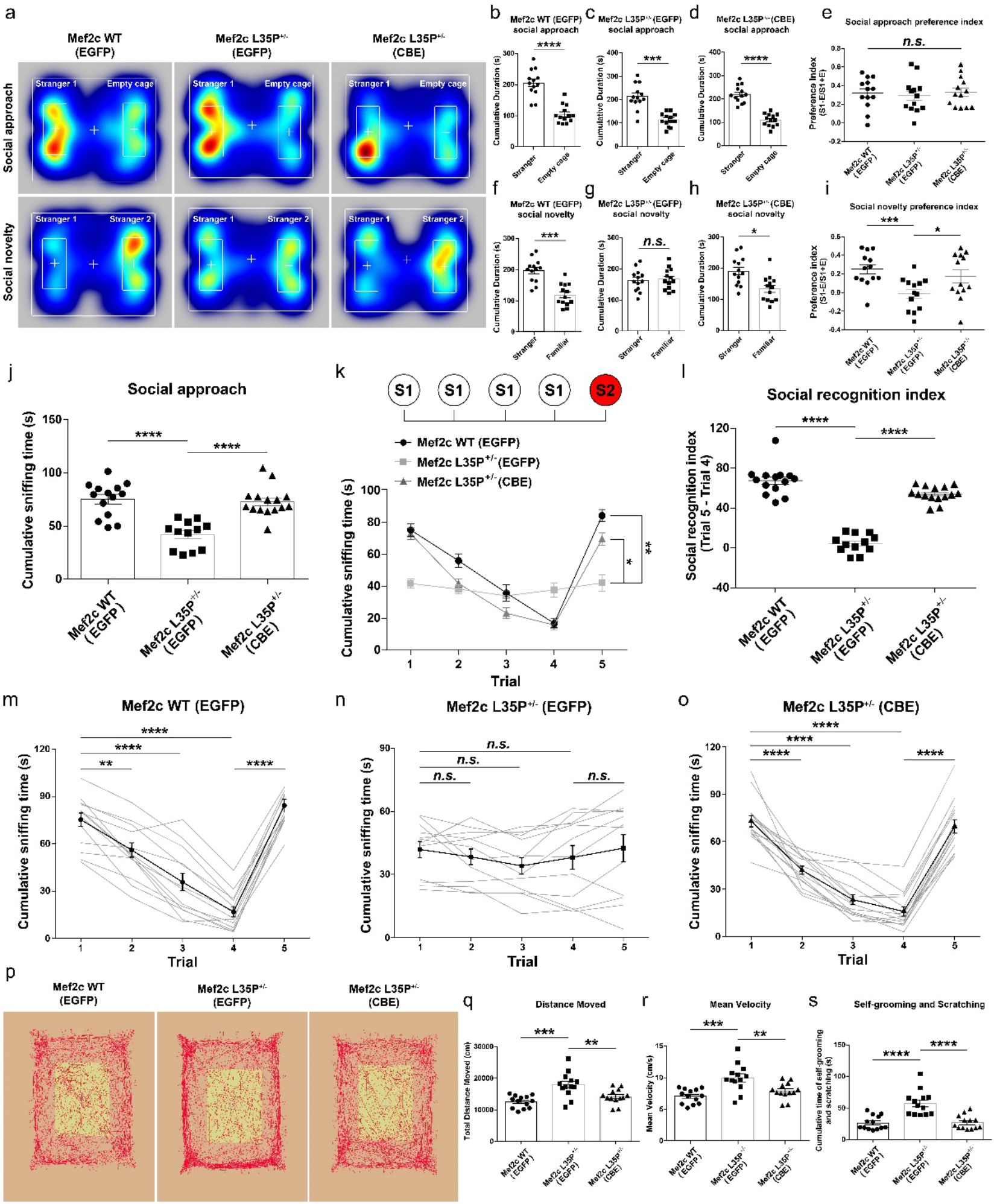
Amelioration of autistic-like behaviors in *Mef2c* L35P^+/−^ mice with CBE gene editing *in vivo*. **a**, Representative locomotion heat maps of *Mef2c* WT mice injection of AAV-EGFP, and *Mef2c* L35P^+/−^ mice injected with either EGFP or CBE in the three-chamber test. Cumulative duration of the social approach session of *Mef2c* WT mice injection of AAV-EGFP (**b**), and *Mef2c* L35P^+/−^ mice injected with either EGFP (**c**) or CBE (**d**) (*n*=13). **e**, the preference index of social approach for three groups. Cumulative duration of the social novelty episode of *Mef2c* WT mice injected with AAV-EGFP (**f**), and *Mef2c* L35P^+/−^ mice injected with either EGFP (**g**) or CBE (**h**) (*n*=13). **i**, the preference index of social novelty test of three groups. **j**, Cumulative sniffing time in the first trial of the social intruder test for *Mef2c* WT mice injected with AAV-EGFP (*n*=14), *Mef2c* L35P^+/−^ mice injected with either EGFP (*n*=12) or CBE (*n*=15). **k**, Cumulative sniffing time in five respective trials of *Mef2c* WT mice injected with AAV-EGFP (*n*=14), *Mef2c* L35P^+/−^ mice injected with either EGFP (*n*=12) or CBE (*n*=15) (two-way ANOVA). **l**, The social recognition index of the social intruder test for three groups (unpaired two-tailed Student’s *t*-test). Mean cumulative sniffing time (black curve) and data for individual mice (gray curve) in the social intruder test for *Mef2c* WT mice injected with AAV-EGFP (**m**, *n*=14), *Mef2c* L35P^+/−^ mice injected with either EGFP (**n**, *n*=12) or CBE (**o**, *n*=15, one-way ANOVA). **p**, Representative locomotion track traces of *Mef2c* WT mice injection of AAV-EGFP and *Mef2c* L35P^+/−^ mice injected with either EGFP or CBE in the open field test. Quantification of total moving distance (**q**) and mean velocity (**r**) of three groups (*n*=13; unpaired two-tailed Student’s *t*-test). **s**, Quantitative analysis of cumulative time spent in self-grooming and scratching of *Mef2c* WT mice injection of AAV-EGFP and *Mef2c* L35P^+/−^ mice injected with either EGFP or CBE (*n*=13 mice per group; unpaired two-tailed Student’s *t*-test). *n* represents mice number. Statistical values represent the mean ± s.e.m. *\*p* < 0.05, *\*\*p* < 0.01, *\*\*\*p* < 0.001, *\*\*\*\*p* < 0.0001.

Finally, we tested if *in vivo* base editing could rescue the hyperactive phenotypes of *Mef2c* L35P^+/−^ mice. In the open field test, we found that *Mef2c* L35P^+/−^ mice treated with dual-AAV CBE displayed similar locomotion activity to WT mice (Fig. 5p-r). We also observed that the increased self-grooming behaviors in *Mef2c* L35P^+/−^ mice were rescued by CBE (Fig. 5s). Taken together, these results indicate that the dual-AAV CBE strategy can potently rescue behavioral deficits in a mouse model of ASD, even if administered postnatally.

## Discussion

Previous reports indicate that *Mef2c^+/−^*mice exhibit behavioral deficits mimicking people with ASD, thus serving as a faithful animal model ^35^. Interestingly, NitroSynapsin, a new dual-action compound similar to the FDA approved drug memantine, is able to rescue the behavioral deficits, E/I imbalance, and histological abnormalities of *Mef2c^+/−^*mice, suggesting that modulating synaptic activity may be able to reverse the autistic-like phenotypes in postnatal animal models of ASD ^35^.

Recently, there are reports showing that application of genome editing tools postnatally in specific brain regions of ASD mouse models, including prefrontal cortex, anterior cingulate cortex and hippocampus, could successfully rescue autistic-like defects, indicating that modulation of synaptic activity via genetic manipulations may also provide a reliable way to regulate animal behaviors during pathological status ^48,49,50^. However, for therapeutic purposes, viral injections into specific regions of the brain of ASD patients may not be ideal approaches so far.

To overcome these obstacles, we have developed an easier way of genome editing targeting neurons in the brain via a BBB-crossing AAV system. We showed that through a newly designed CBE system and intein-mediated split strategy, CRISPR-based CBE could be successfully delivered into the brain via dual AAVs and performed base editing effectively. Though the AAV-PHP.eB vector usually preferentially infected neurons in cortical and hippocampal regions, we observed that the protein level of MEF2C in the specific brain regions and autistic-like behaviors of MEF2C-L35P mice, including abnormal social interaction and hyperactivity, were both largely rescued. These results establish the framework of using genome base editing tools *in vivo* in the brain to correct genetic mutations and provide potential therapeutic approaches for people with ASD. Because we were able to reverse phenotypes even postnatally, it gives hope for the development of therapeutic approaches for adolescents and adults living with ASD.

## Lead Contact

Further information and requests for resources and reagents should be directed to and will be fulfilled by the Lead Contact, Zilong Qiu.

## Ethics approval and consent to participate

The genetic information collected from the ASD patient is approved by the Ethic Committee of Xinhua hospital, Shanghai Jiao Tong University School of Medicine (XHEC-C-2019-076).

## Availability of data and materials

The datasets used and/or analyzed during the current study are available from the lead contact on reasonable request.

## Acknowledgements

We thank the families for their participation in this study. We thank Dr. Aaron Gitler for critical comments of the manuscript. This work was supported by grants from the NSFC Grants (#31625013, #81941015, #82021001); Strategic Priority Research Program of the Chinese Academy of Sciences (XDB32060202); Program of Shanghai Academic Research Leader, the Open Large Infrastructure Research of Chinese Academy of Sciences, the Shanghai Municipal Science and Technology Major Project (#2018SHZDZX05) and the Guangdong Key Project (2018B030335001). Z.Q. is supported by GuangCi Professorship Program of Ruijin Hospital Shanghai Jiao Tong University School of Medicine. This work was supported by grants from National Key R&D Program of China (2019YFA0111000), Natural Science Foundation of Shanghai (20ZR1403100), Shanghai Municipal Science and Technology (20JC1419500) to T.L.C., National Natural Science Foundation of China (#31600826) to T.L.C.

## References

1 Colvert, E. et al., Heritability of Autism Spectrum Disorder in a UK Population-Based Twin Sample. JAMA PSYCHIAT 72 415 (2015).

2 Geschwind, D. H. & Flint, J., Genetics and genomics of psychiatric disease. SCIENCE 349 1489 (2015).

3 Skuse, D. H., Mandy, W. P. & Scourfield, J., Measuring autistic traits: heritability, reliability and validity of the Social and Communication Disorders Checklist. Br J Psychiatry 187 568 (2005).

4 Iossifov, I. et al., De novo gene disruptions in children on the autistic spectrum. NEURON 74 285 (2012).

5 Sanders, S. J. et al., De novo mutations revealed by whole-exome sequencing are strongly associated with autism. NATURE 485 237 (2012).

6 Parikshak, N. N. et al., Integrative functional genomic analyses implicate specific molecular pathways and circuits in autism. CELL 155 1008 (2013).

7 Gilissen, C. et al., Genome sequencing identifies major causes of severe intellectual disability. NATURE 511 344 (2014).

8 Leifer, D. et al., MEF2C, a MADS/MEF2-family transcription factor expressed in a laminar distribution in cerebral cortex. Proc Natl Acad Sci U S A 90 1546 (1993).

9 Okamoto, S., Krainc, D., Sherman, K. & Lipton, S. A., Antiapoptotic role of the p38 mitogen-activated protein kinase-myocyte enhancer factor 2 transcription factor pathway during neuronal differentiation. Proc Natl Acad Sci U S A 97 7561 (2000).

10 Shalizi, A. et al., A calcium-regulated MEF2 sumoylation switch controls postsynaptic differentiation. SCIENCE 311 1012 (2006).

11 Flavell, S. W. et al., Activity-dependent regulation of MEF2 transcription factors suppresses excitatory synapse number. SCIENCE 311 1008 (2006).

12 Harrington, A. J. et al., MEF2C regulates cortical inhibitory and excitatory synapses and behaviors relevant to neurodevelopmental disorders. ELIFE 5(2016).

13 Harrington, A. J. et al., MEF2C Hypofunction in Neuronal and Neuroimmune Populations Produces MEF2C Haploinsufficiency Syndrome-like Behaviors in Mice. Biol Psychiatry 88 488 (2020).

14 Barbosa, A. C. et al., MEF2C, a transcription factor that facilitates learning and memory by negative regulation of synapse numbers and function. Proc Natl Acad Sci U S A 105 9391 (2008).

15 Engels, H. et al., A novel microdeletion syndrome involving 5q14.3-q15: clinical and molecular cytogenetic characterization of three patients. EUR J HUM GENET 17 1592 (2009).

16 Le Meur, N. et al., MEF2C haploinsufficiency caused by either microdeletion of the 5q14.3 region or mutation is responsible for severe mental retardation with stereotypic movements, epilepsy and/or cerebral malformations. J MED GENET 47 22 (2010).

17 Paciorkowski, A. R. et al., MEF2C Haploinsufficiency features consistent hyperkinesis, variable epilepsy, and has a role in dorsal and ventral neuronal developmental pathways. NEUROGENETICS 14 99 (2013).

18 Hsu, P. D., Lander, E. S. & Zhang, F., Development and applications of CRISPR-Cas9 for genome engineering. CELL 157 1262 (2014).

19 Doudna, J. A. & Charpentier, E., Genome editing. The new frontier of genome engineering with CRISPR-Cas9. SCIENCE 346 1258096 (2014).

20 Cai, Y. et al., In vivo genome editing rescues photoreceptor degeneration via a Cas9/RecA-mediated homology-directed repair pathway. SCI ADV 5 v3335 (2019).

21 Gaudelli, N. M. et al., Programmable base editing of A*T to G*C in genomic DNA without DNA cleavage. NATURE 551 464 (2017).

22 Komor, A. C., Kim, Y. B., Packer, M. S., Zuris, J. A. & Liu, D. R., Programmable editing of a target base in genomic DNA without double-stranded DNA cleavage. NATURE 533 420 (2016).

23 Rees, H. A. & Liu, D. R., Base editing: precision chemistry on the genome and transcriptome of living cells. NAT REV GENET 19 770 (2018).

24 Koblan, L. W. et al., In vivo base editing rescues Hutchinson-Gilford progeria syndrome in mice. NATURE 589 608 (2021).

25 Newby, G. A. & Liu, D. R., In vivo somatic cell base editing and prime editing. MOL THER 29 3107 (2021).

26 Cheng, T. L. et al., Expanding C-T base editing toolkit with diversified cytidine deaminases. NAT COMMUN 10 3612 (2019).

27 Duan, Y. et al., Brain-wide Cas9-mediated cleavage of a gene causing familial Alzheimer’s disease alleviates amyloid-related pathologies in mice. NAT BIOMED ENG (2021).

28 Wang, J. et al., Novel MEF2C point mutations in Chinese patients with Rett (-like) syndrome or non-syndromic intellectual disability: insights into genotype-phenotype correlation. BMC MED GENET 19 191 (2018).

29 Le Meur, N. et al., MEF2C haploinsufficiency caused by either microdeletion of the 5q14.3 region or mutation is responsible for severe mental retardation with stereotypic movements, epilepsy and/or cerebral malformations. J MED GENET 47 22 (2010).

30 Zweier, M. et al., Mutations in MEF2C from the 5q14.3q15 microdeletion syndrome region are a frequent cause of severe mental retardation and diminish MECP2 and CDKL5 expression. HUM MUTAT 31 722 (2010).

31 Bienvenu, T., Diebold, B., Chelly, J. & Isidor, B., Refining the phenotype associated with MEF2C point mutations. NEUROGENETICS 14 71 (2013).

32 Srivastava, S. et al., Clinical whole exome sequencing in child neurology practice. ANN NEUROL 76 473 (2014).

33 Vrecar, I. et al., Further Clinical Delineation of the MEF2C Haploinsufficiency Syndrome: Report on New Cases and Literature Review of Severe Neurodevelopmental Disorders Presenting with Seizures, Absent Speech, and Involuntary Movements. J Pediatr Genet 6 129 (2017).

34 Li, C. et al., MKRN3-mediated ubiquitination of Poly(A)-binding proteins modulates the stability and translation of GNRH1 mRNA in mammalian puberty. NUCLEIC ACIDS RES 49 3796 (2021).

35 Tu, S. et al., NitroSynapsin therapy for a mouse MEF2C haploinsufficiency model of human autism. NAT COMMUN 8 1488 (2017).

36 Lyons, G. E., Micales, B. K., Schwarz, J., Martin, J. F. & Olson, E. N., Expression of mef2 genes in the mouse central nervous system suggests a role in neuronal maturation. J NEUROSCI 15 5727 (1995).

37 Durand, S. et al., NMDA receptor regulation prevents regression of visual cortical function in the absence of Mecp2. NEURON 76 1078 (2012).

38 He, L. J. et al., Conditional deletion of Mecp2 in parvalbumin-expressing GABAergic cells results in the absence of critical period plasticity. NAT COMMUN 5 5036 (2014).

39 Tong, D. L. et al., The critical role of ASD-related gene CNTNAP3 in regulating synaptic development and social behavior in mice. NEUROBIOL DIS 130 104486 (2019).

40 Foss-Feig, J. H. et al., Searching for Cross-Diagnostic Convergence: Neural Mechanisms Governing Excitation and Inhibition Balance in Schizophrenia and Autism Spectrum Disorders. Biol Psychiatry 81 848 (2017).

41 Sohal, V. S. & Rubenstein, J., Excitation-inhibition balance as a framework for investigating mechanisms in neuropsychiatric disorders. Mol Psychiatry 24 1248 (2019).

42 Walton, R. T., Christie, K. A., Whittaker, M. N. & Kleinstiver, B. P., Unconstrained genome targeting with near-PAMless engineered CRISPR-Cas9 variants. SCIENCE 368 290 (2020).

43 Zhou, C. et al., Off-target RNA mutation induced by DNA base editing and its elimination by mutagenesis. NATURE 571 275 (2019).

44 Wang, X. et al., Efficient base editing in methylated regions with a human APOBEC3A-Cas9 fusion. NAT BIOTECHNOL 36 946 (2018).

45 Liu, Y. et al., A Cas-embedding strategy for minimizing off-target effects of DNA base editors. NAT COMMUN 11 6073 (2020).

46 Zettler, J., Schutz, V. & Mootz, H. D., The naturally split Npu DnaE intein exhibits an extraordinarily high rate in the protein trans-splicing reaction. FEBS LETT 583 909 (2009).

47 Yang, K. et al., SENP1 in the retrosplenial agranular cortex regulates core autistic-like symptoms in mice. CELL REP 37 109939 (2021).

48 Guo, B. et al., Anterior cingulate cortex dysfunction underlies social deficits in Shank3 mutant mice. NAT NEUROSCI 22 1223 (2019).

49 Yu, B. et al., Reversal of Social Recognition Deficit in Adult Mice with MECP2 Duplication via Normalization of MeCP2 in the Medial Prefrontal Cortex. NEUROSCI BULL 36 570 (2020).

50 Sun, L. et al., Visualization and correction of social abnormalities-associated neural ensembles in adult MECP2 duplication mice. SCI BULL 65 1192 (2020).

