## Supplementary figures and methods for "Whole-brain *in vivo* base editing reverses autistic-like behaviors in mice"

**Supplementary Figures 1-8**

**Supplementary Figure Legends**

**Supplementary Materials and Methods**

**Figure S1-S8**

### Supplementary Figure 1

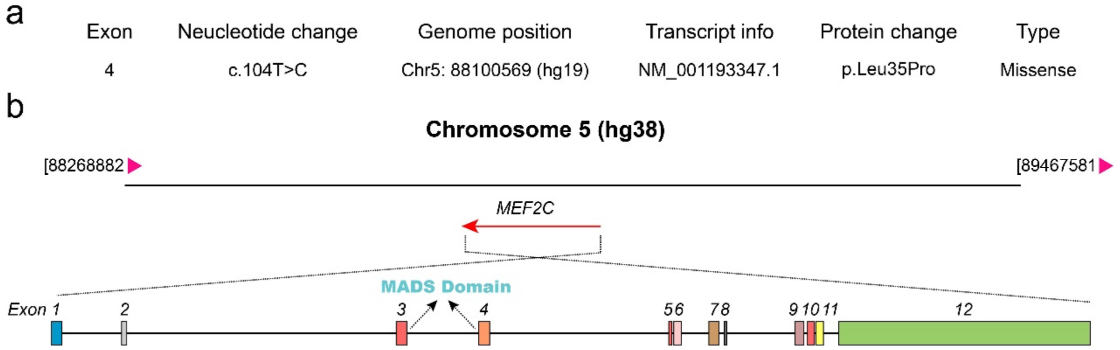


**Supplementary Figure 1. The annotation of *de novo* single nucleotide mutation in human *MEF2C* gene.**

**a**, Annotation of the genomic and transcript information of L35P. (based on the hg19 database). **b**, Schematic diagram of the location of human *MEF2C* gene in chromosome and the genomic profile of MEF2C alternative splicing isoform 3 (NM_001193347.1).

### Supplementary Figure 2

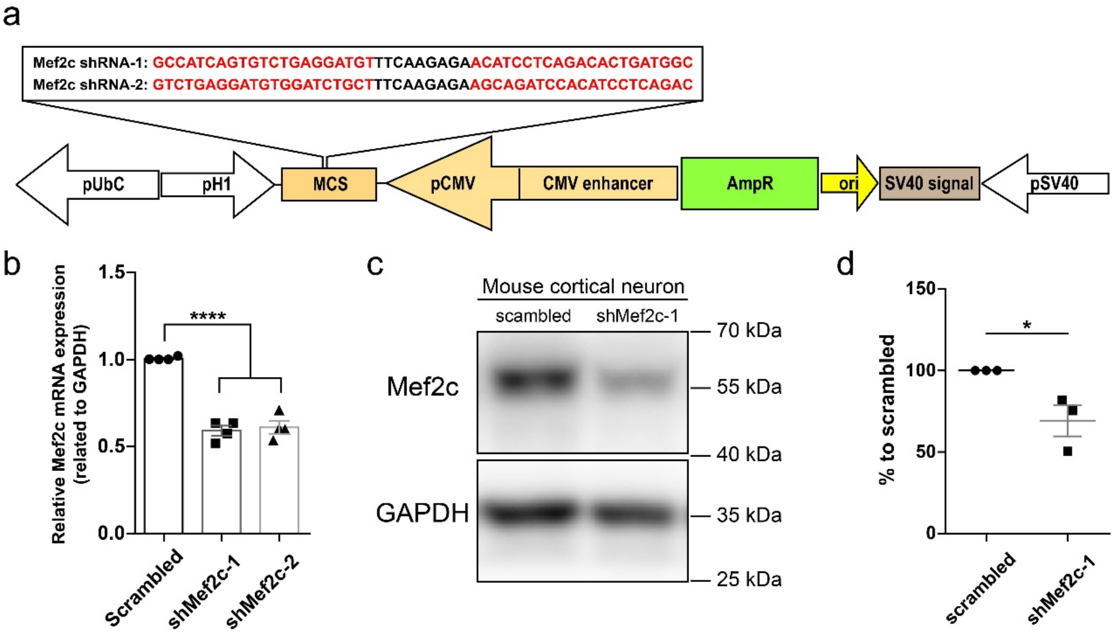


**Supplementary Figure 2. Design a of mouse *Mef2c* shRNA constructs and validation of shRNA knock-down efficiency.**

**a**, Schematic illustration of engineering of two different *Mef2c* shRNA designed based on BLOCK-iT™ RNAi Designer (http://rnaidesigner.thermofisher.com/rnaiexpress/). **b**, Quantification of *Mef2c* mRNA level of cultured primary cortical neurons infected with lentiviral vectors expressing scrambled or two designed *Mef2c* shRNA v ($n$=4 per group, one-way ANOVA). **c**, Representative western blotting of cultured primary cortical neurons infected with lentiviral vectors expressing scrambled or *Mef2c* shRNA-1 for 72 h. Cells were harvested at DIV5. GAPDH was used as internal control. **d**, Quantitative analysis of relative *Mef2c* protein expression. Neurons infected with scrambled shRNA were normalized to 100% ($n$=3, unpaired two-tailed Student's $t$-test). $n$ is biological repeat numbers of independent experiments. Statistical values represent the mean ± s.e.m. $*p < 0.05$, $****p < 0.0001$.

**Supplementary Figure 3**

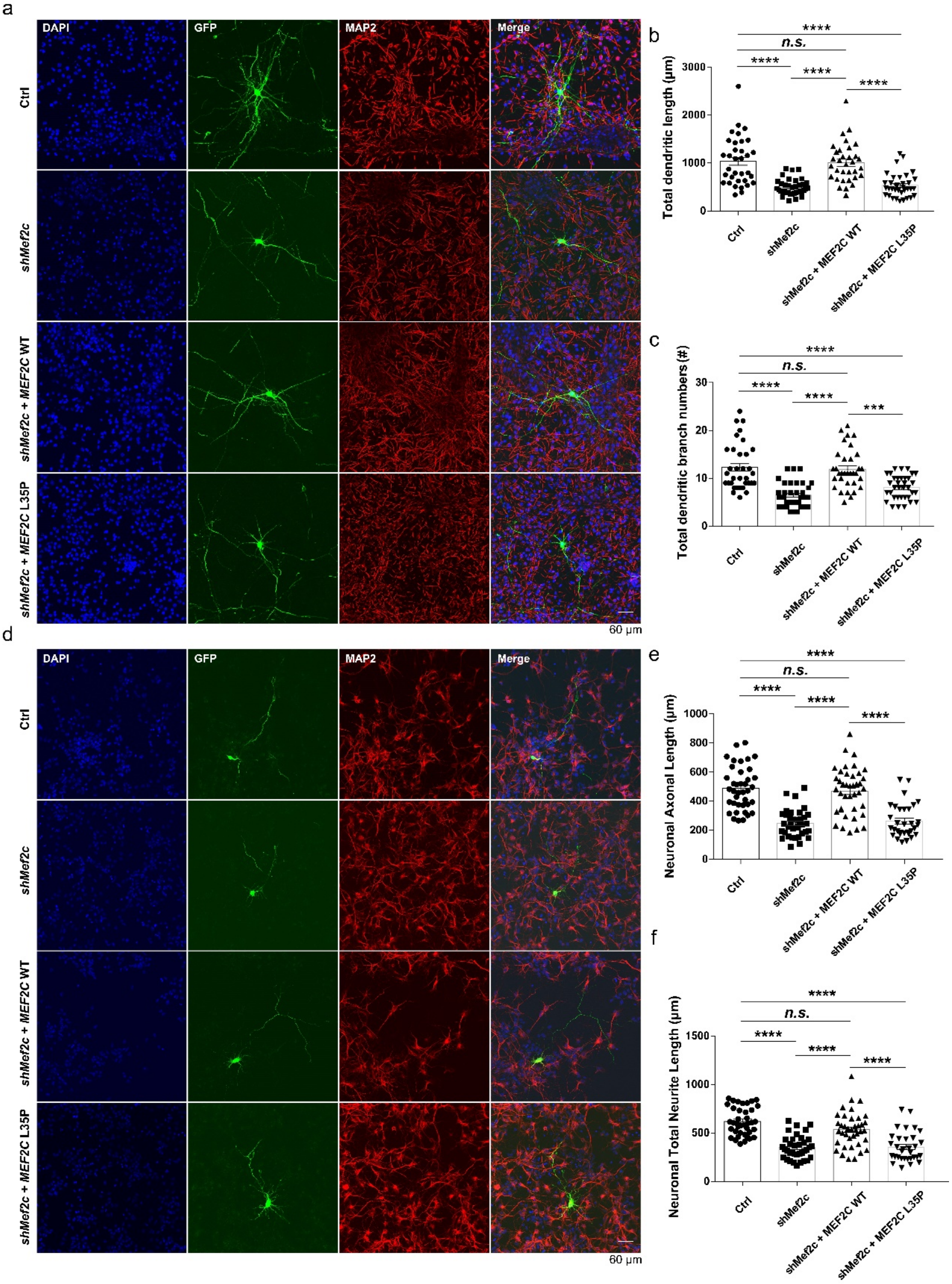


**Supplementary Figure 3. Impaired neuronal dendritic and axonal morphological development in *Mef2c* knock down neurons could be rescued by MEF2C -WT, but not MEF2C-L35P.**

**a**, Representative image of immunofluorescent staining for GFP (green), MAP2 (red) and DAPI (blue) of cultured E14.5 mouse cortical neurons. Mouse cortical neurons were transfected with DsRed shRNA (Ctrl, *n*=39), shMef2c (*n*=36), shMef2c with human MEF2C-WT (*n*=40) or MEF2C-L35P (*n*=32) respectively at DIV1 via Lipofectamine 3000. Neuronal cells were immunofluorescent stained at DIV10. Scale bars, 60μm. Quantitative analysis of neuronal total dendritic length (**b**) and branch numbers (**c**) of neurons transfected with Ctrl, shMef2c, shMef2c+MEF2C-WT and shMef2c+MEF2C-L35P. **d**, Representative immunofluorescent staining images for GFP (green), MAP2 (red) and DAPI (blue) of cultured E14.5 mouse cortical neurons. Neuronal cells were transfected with DsRed shRNA (Ctrl, *n*=34), shMef2c (*n*=37), shMef2c with human MEF2C-WT (*n*=33) or MEF2C-L35P (*n*=34) respectively at DIV1. Neuronal cells were immunofluorescent stained at DIV4. Scale bars, 60μm. Quantitative analysis of neuronal axonal (**e**) and total neurite length (**f**) of neurons transfected with Ctrl, shMef2c, shMef2c+MEF2C-WT and shMef2c+MEF2C-L35P. *n* is biological repeat numbers of independent experiments. Statistical values represent the mean ± s.e.m. $^{***}p < 0.001$, $^{****}p < 0.0001$, One-way ANOVA.

### Supplementary Figure 4

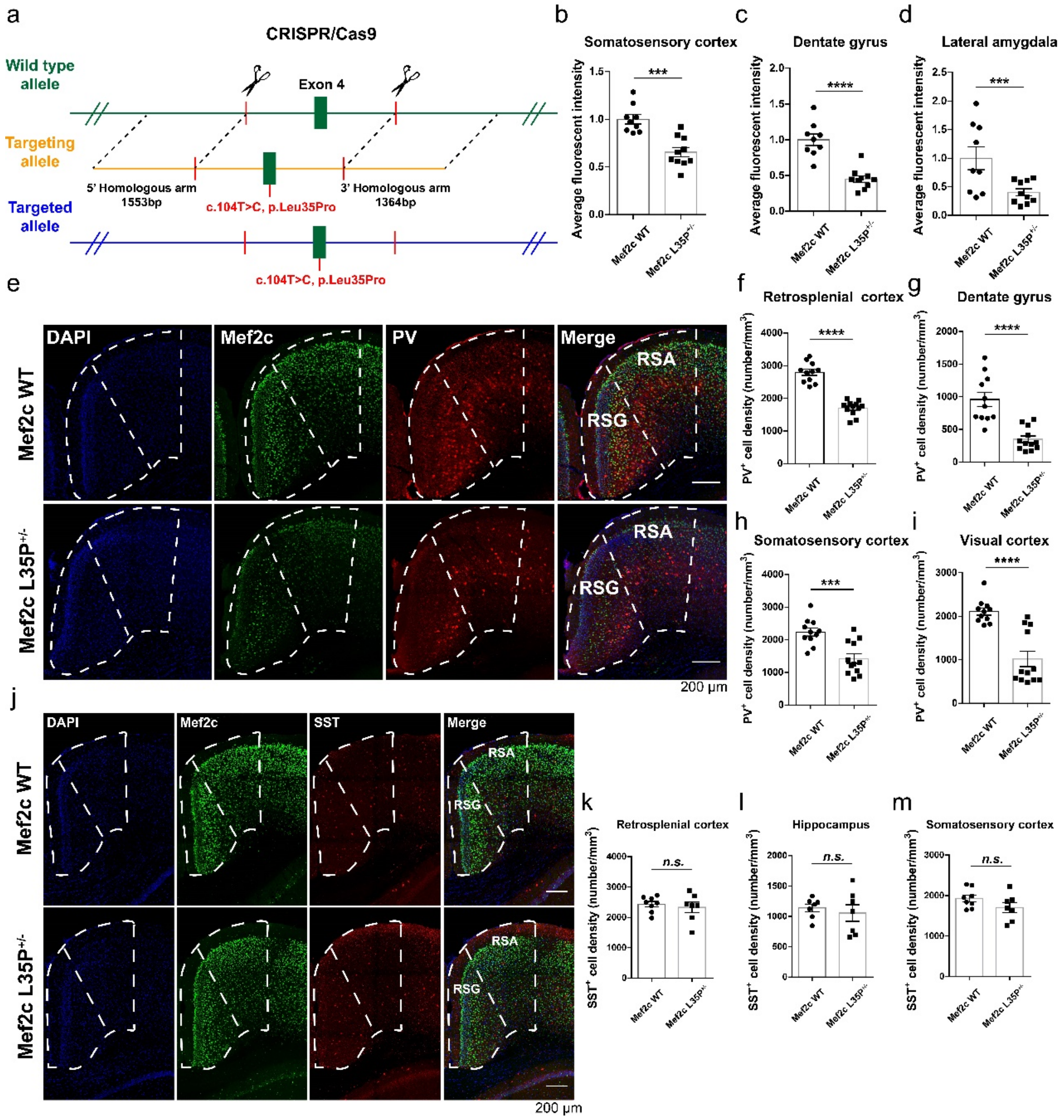


**Supplementary Figure 4. Decreased protein expression and aberrant inhibitory interneuron development in *Mef2c* L35P$^{+/-}$ mice.**

**a**, Design and construction of *Mef2c* L35P knock-in mice by CRISPR/Cas9 mediated homologous recombination. Quantitative analysis of average fluorescent intensity of Mef2c in **Figure 2d**: somatosensory cortex (**b**), dentate gyrus (**c**) and lateral amygdala (**d**) of *Mef2c* WT (*n*=9 brain slices from 3 mice) and L35P$^{+/-}$ mice (*n*=10 brain slices from 3 mice). **e**, Representative images of immunohistochemical staining for PV (red), Mef2c (green) and DAPI (blue) in the retrosplenial cortex (white dotted box indicates corresponding brain region) of *Mef2c* WT and L35P$^{+/-}$ mice. Scale bars, 200μm. Quantitative analysis of PV-positive neuronal cell density in the retrosplenial cortex (**f**), dentate gyrus (**g**), somatosensory cortex (**h**) and visual cortex (**i**) of *Mef2c* WT ($n$=11 brain slices from 4 mice) and L35P$^{+/-}$ mice ($n$=12 brain slices from 4 mice). **j**, Representative images of immunohistochemical staining for SST (red), Mef2c (green) and DAPI (blue) in the retrosplenial cortex (White dotted box indicates corresponding brain region) of *Mef2c* WT and L35$^{+/-}$ mice. Scale bars, 200μm. Quantification of SST-positive neuronal cell density in the retrosplenial cortex (**k**), hippocampus (**l**) and somatosensory cortex (**m**) of *Mef2c* WT ($n$=8 brain slices from 3 mice) and L35P$^{+/-}$ mice ($n$=7 brain slices from 3 mice). Statistical values represent the mean ± s.e.m. ***$p$ < 0.001, ****$p$ < 0.0001, unpaired two-tailed Student's $t$-test.

### Supplementary Figure 5

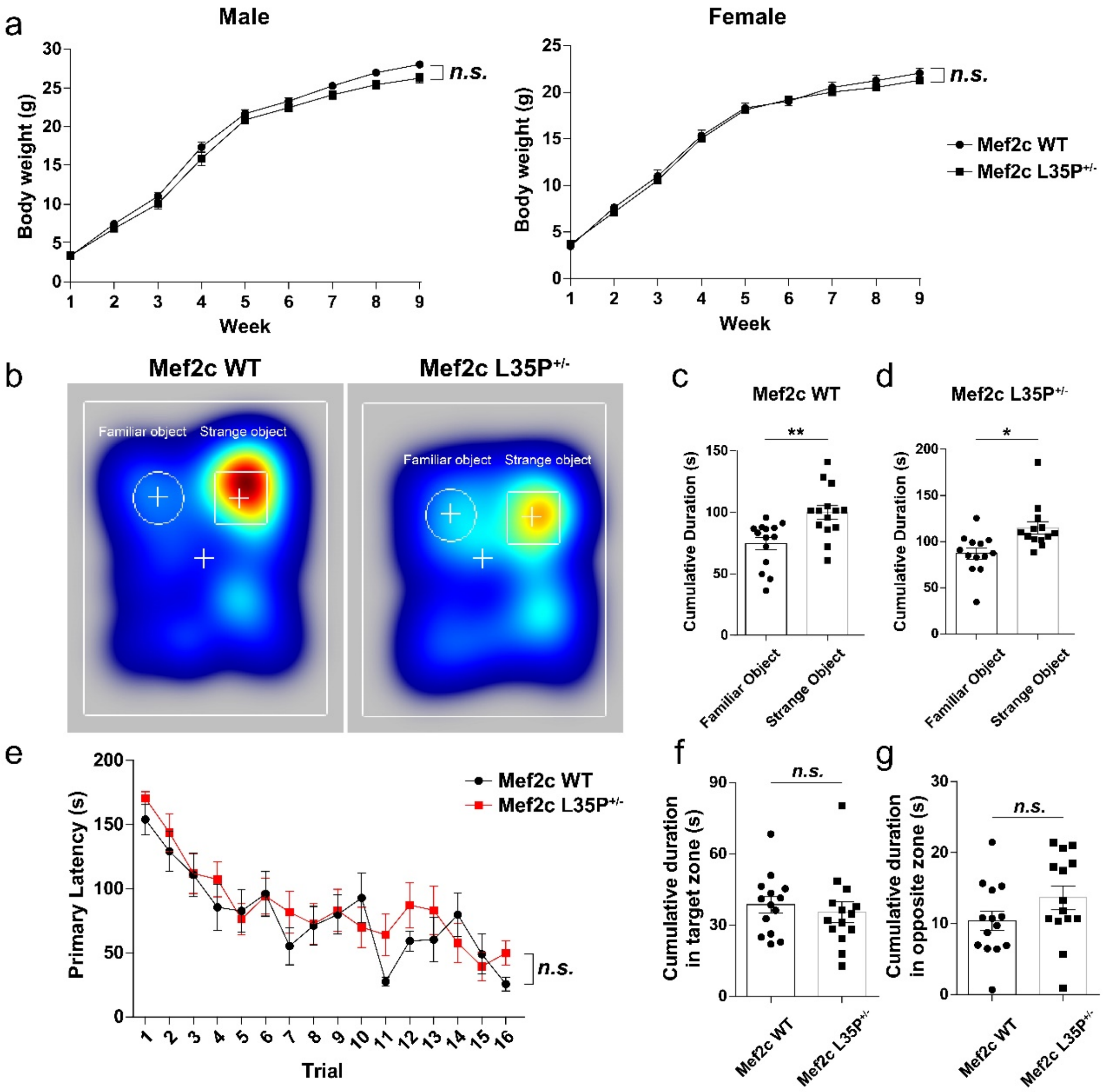


**Supplementary Figure 5. General development and cognitive ability of *Mef2c***

**L35P$^{+/-}$ mice.**

**a**, Body weights of *Mef2c* WT and L35P$^{+/-}$ mice of both sex from postnatal one week to nine weeks. (WT, $n$=29 male mice, $n$=17 female mice; L35P$^{+/-}$, $n$=17 male mice, $n$=21 female mice; two-way ANOVA). **b**, Representative heat maps of *Mef2c* WT and L35P$^{+/-}$ mice in the novel object recognition task (while circle indicates familiar object; white box indicates stranger object). Quantification of cumulative sniffing time of *Mef2c* WT (**c**) and L35P$^{+/-}$ mice (**d**) in the task (WT, $n$=14; L35P$^{+/-}$, $n$=13; unpaired two-tailed Student's $t$-test). **e**, Quantitative analysis of primary latency spent in locating the target hole of Mef2c WT and L35P+/- mice in the Barnes maze ($n$=14 per group; two-way ANOVA). Quantification of cumulative duration exploring in the target (**f**) and opposite zone (**g**) of *Mef2c* WT and L35P$^{+/-}$ mice in the apparatus, ($n$=14 per group; unpaired two-tailed Student's $t$-test). $n$ represents mice number. Statistical values represent the mean ± s.e.m. *$p < 0.05$, **$p < 0.01$.

### Supplementary Figure 6

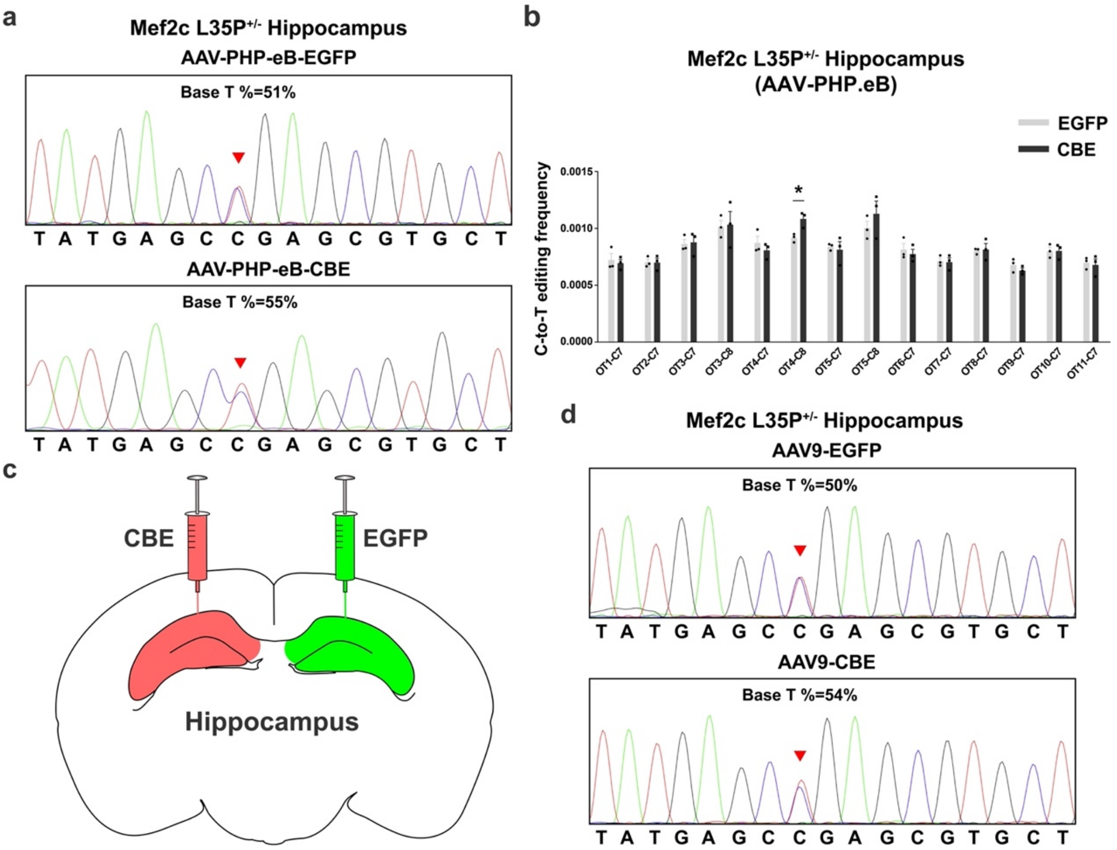

**Supplementary Figure 6. Identification of *in vivo* base editing efficiency and off-target analysis in hippocampus of *Mef2c* L35P$^{+/-}$ mice.**

**a**, Representative sanger sequencing results of hippocampus tissues extracted from *Mef2c* L35P$^{+/-}$ mice treated with either AAV-PHP.eB delivered EGFP or CBE (red arrowhead indicate gene loci treated with or without gene editing; Proportion of thymine (T) base pair after CBE-dependent gene repair was 55%, while treated with EGFP was 51%). **b**, Quantitative analysis of off-target editing efficiency (C-to-T conversion) in *Mef2c* L35P$^{+/-}$ mice hippocampi tissues injected with either AAV-PHP.eB mediated EGFP or CBE ($n$=3). **c**, Schematical illustration of local stereotactic injection method in the whole hippocampus of *Mef2c* L35P$^{+/-}$ mice. Unilateral hippocampus was injected with AAV9 delivered EGFP while contralateral hippocampus was treated with CBE. **d**, Representative sanger sequencing results of *Mef2c* L35P$^{+/-}$ mice hippocampus injected with either AAV9 mediated EGFP or CBE (red arrowhead indicate gene loci treated with or without gene editing; Proportion of thymine (T) base pair after CBE-dependent gene therapy was 54%, while injected with EGFP was 50%). The percentage of T (c.104T) is obtained by calculating T base ratio (T /(C+T) * 100%). $n$ represents mice number. Statistical values represent the mean ± s.e.m. *$p < 0.05$, unpaired two-tailed Student's $t$-test.

### Supplementary Figure 7

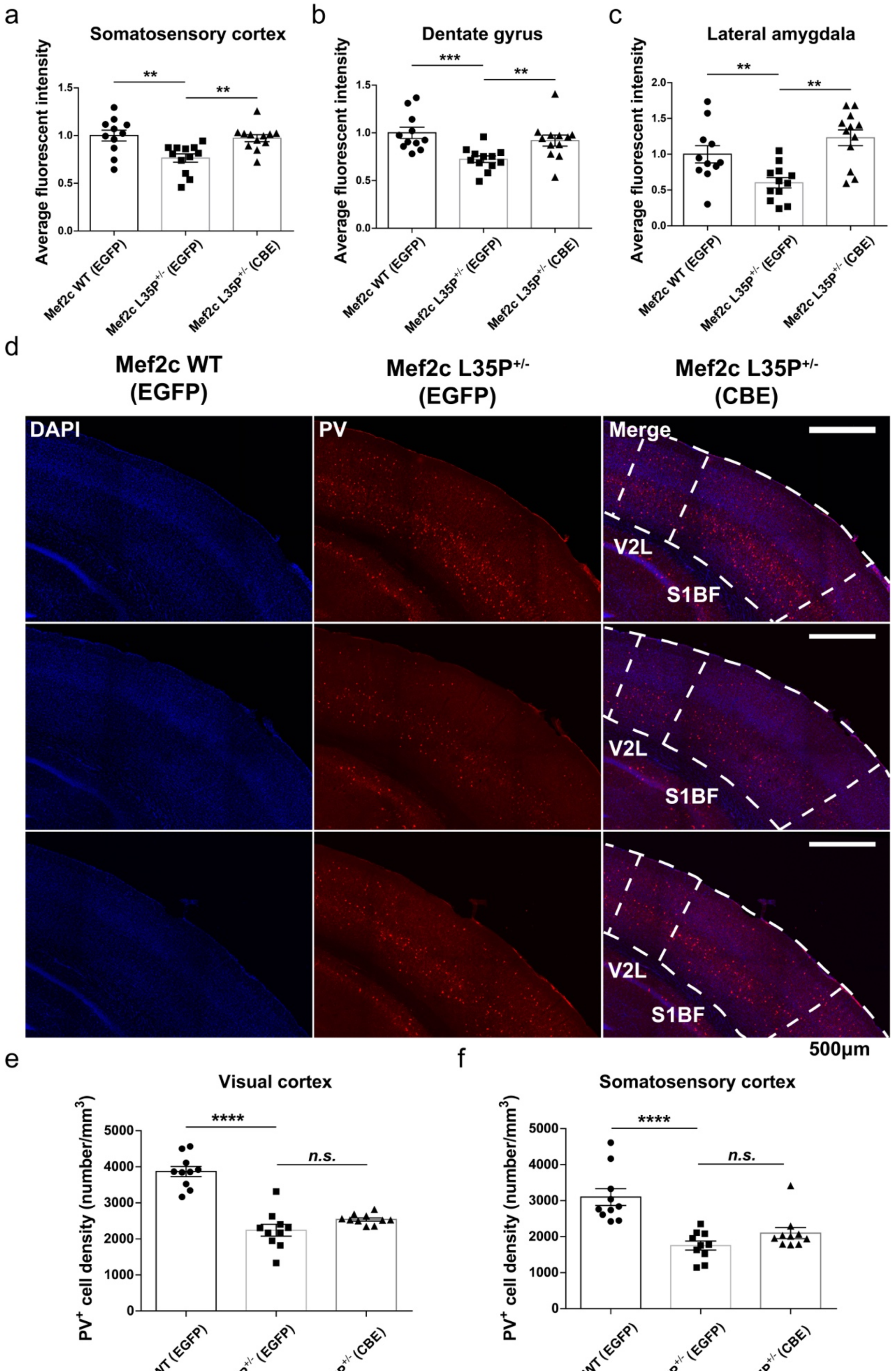

**Supplementary Figure 7.** ***In vivo*** **CBE gene editing did not restore decreased PV-positive inhibitory interneuron in visual and somatosensory cortex of** ***Mef2c*** **L35P$^{+/-}$ mice**

Quantification of average fluorescent intensity of Mef2c in the somatosensory cortex (**a**), dentate gyrus (**b**) and lateral amygdala (**c**) of *Mef2c* WT EGFP (*n*=11 brain slices from 4 mice) L35P$^{+/-}$ EGFP (*n*=12 brain slices from 4 mice) and L35P$^{+/-}$ CBE mice, which was shown in **Figure 4h** (*n*=12 brain slices from 3 mice). **d**, Representative images of immunohistochemical staining for PV (red) and DAPI (blue) in the visual cortex and somatosensory cortex (White dotted box indicates corresponding brain region) of *Mef2c* WT mice injected with AAV-EGFP, *Mef2c* L35P$^{+/-}$ mice injected with either EGFP or CBE. Scale bars, 500µm. Quantitative analysis of PV-positive neuronal cell density in the visual (**e**) and somatosensory cortex (**f**) of three groups (*n*=10 brain slices from 4 mice per group). Statistical values represent the mean ± s.e.m. $^{**}p < 0.01$, $^{***}p < 0.001$, $^{****}p < 0.0001$, unpaired two-tailed Student's *t*-test.

### Supplementary Figure 8

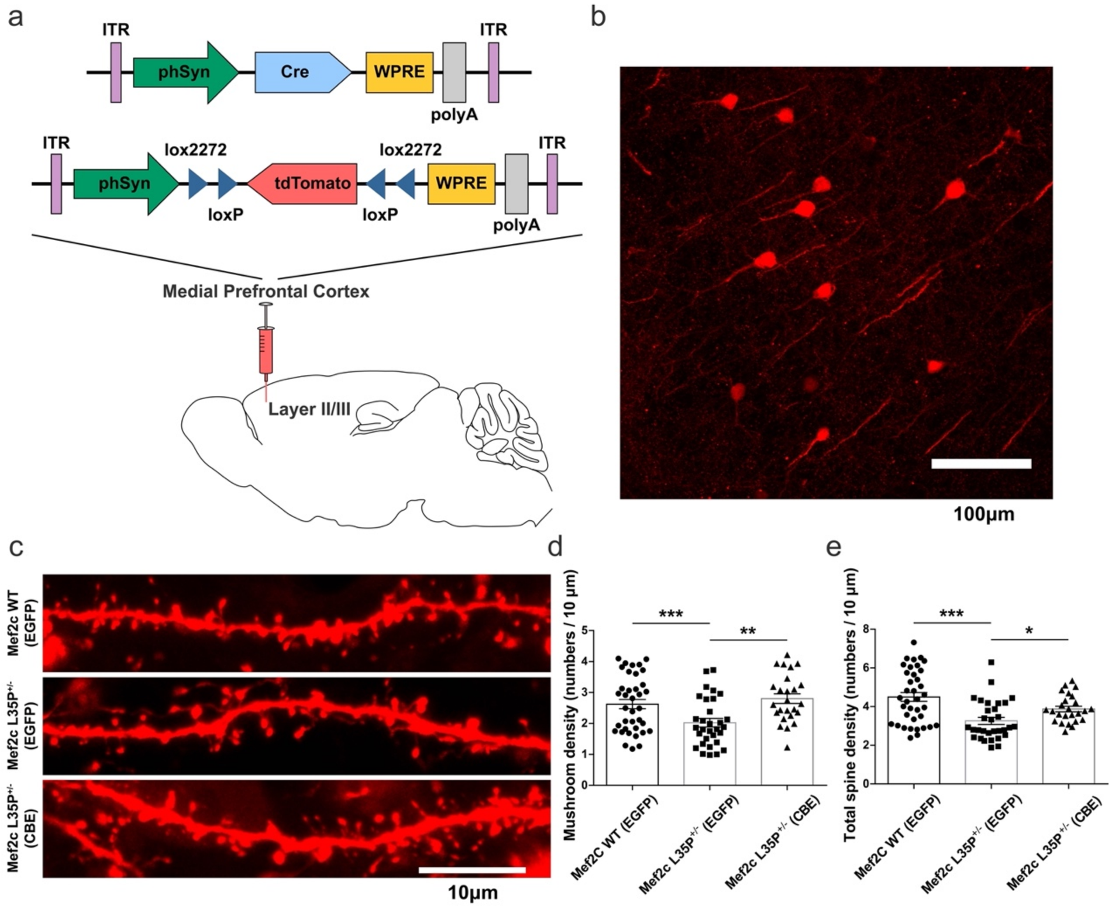


**Supplementary Figure 8. Restoration of mushroom shaped and total spine density in the medial prefrontal cortex layer II/III in *Mef2c* L35P$^{+/-}$ mice after *in vivo* CBE gene editing.**

**a**, Schematic illustration of local stereotactic injection method applied to sparsely label cortical pyramidal neurons in the mPFC layer II/III of *Mef2c* WT mice injected with AAV-EGFP, *Mef2c* L35P$^{+/-}$ mice injected with either EGFP or CBE. **b**, Representative images of cortical pyramidal neurons after sparse labelling in the mPFC layer II/III. Scale bars, 100μm. **c**, Representative immunofluorescent images of primary cortical neuronal dendrites obtained from *Mef2c* WT mice treated with AAV-EGFP, *Mef2c* L35P$^{+/-}$ mice injected with either EGFP or CBE. Scale bars, 10μm. Quantitative analysis of mushroom (**d**) and total spine density (**e**) of cortical pyramidal primary dendrites in the mPFC layer II/III of *Mef2c* WT mice injected with AAV-EGFP (*n*=37 dendrites from 3 mice), *Mef2c* L35P$^{+/-}$ mice injected with EGFP (*n*=31 dendrites from 3 mice) and *Mef2c* L35P$^{+/-}$ mice injected with CBE (*n*=25 dendrites from 3 mice).

Statistical values represent the mean ± s.e.m. *$p < 0.05$, **$p < 0.01$, ***$p < 0.001$, unpaired two-tailed Student's *t*-test.

**Supplementary Materials and Methods**

**Discovery of *MEF2C* rare *de novo* SNV and clinical information on the patient harboring *MEF2C* Leu35Pro.**

Previously, we collected and sequenced blood genomes acquired from one aged two years and half boy with ASD related phenotypes as well as his parents with Xinhua hospital affiliated to Shanghai Jiaotong University School of Medicine and the patient clinically diagnosed with Rett-like syndrome was based on the definition of Diagnostic and Statistical Manual of Mental Disorders, Fifth Edition. Additionally, on the aspects of bioinformatics analysis of the *MEF2C* variants discovered and identified in this article, the longest of *MEF2C* transcript isoform was used as a reference sequence (NM_001193347.1, NP_001180276.1). According to the clinical diagnosis of Xinhua hospital, the patient with autosome *MEF2C* rare *de novo* SNV, Leu35Pro exhibited neuronal developmental retardation, poor eye contact, motor abnormality and involuntary hand movement.

**Engineer and construction of plasmids.**

Human *MEF2C* WT transcript 3 (NM_001193347.1) CDS sequence, which was synthesized in Genescript corporation, was built into empty FUGW-EGFP driven by UbC promoter (Addgene plasmid 81018; using for HEK293 cell transfection) and pCAGGS driven by CAG promoter (modified by our lab; using for neuronal cell transfection) vector. On the aspect of the construction of *MEF2C* L35P (L35F, L35I, L35A) vectors, we adopted KOD-Plus- Mutagenesis Kit (TOYOBO Bio-Technology, CO., LTD.) to reproduce T to C (corresponding replacement) conversion on the position of *MEF2C* WT CDS, which was an inverse PCR (iPCR)-based site-directed mutagenesis kit using KOD-plus DNA polymerase (No. KOD-201) as a high-fidelity PCR enzyme. To be mentioned, IRES sequence was inserted into the location between human *MEF2C* WT CDS and EGFP, and additionally FLAG tag was inserted into the plasmid following *MEF2C* WT CDS. In addition, two different shRNA were inserted into FUGW-H1 vector (Addgene plasmid 25870) for knock-down assays. Besides, the empty FUGW-EGFP vector was co-transfected into cultured primary mouse cortical neurons for enhancing the intensity of GFP fluorescence. What needed to be mentioned was that these shRNA sequence we designed was located at the non-homologous region of mouse *Mef2c* transcript 3 through aligning to human *MEFC* transcript 3, suggesting that these two designed shRNA were not available to target to *MEF2C* mRNA to achieve the purpose of knock-down. All FUGW, FUGW-H1, pCAGGS plasmids were packaged into lentivirus for improving transfection efficiency in cultured mouse neurons.

For base editing efficiency of APOBEC3A-CBE system *in vitro*, *Mef2c* sgRNA was cloned into pU6-sgRNA-EF1α-UGI-T2A-mCherry plasmid linearized with BsaI, and paired oligonucleotides were synthesized, annealed, and inserted for sgRNA expression vectors construction. Besides, pCBH-SpG-1249-APOBEC3A$^{Y130F}$-T2A-EGFP plasmid was engineered with APOBEC3A inserted into SpG on the position of 1249amino acid (PX461, Addgene plasmid 48140) [1].

For *Mef2c* gene correction in *Mef2c* L35P$^{+/-}$ mice by means of delivery of APOBEC3A-CBE system, AAV-hSyn-1-1248-pU6-sgRNA plasmid was linearized with BsmBI, and paired oligonucleotides were inserted for sgRNA expression. AAV-hSyn-1249-APOBEC3A$^{Y130F}$-1368-UGI plasmid was engineered of AAV-hSyn-1249-SpG plasmid linearized with SpeI and BsrGI, and APOBEC3A$^{Y130F}$-UGI component with same enzymes, and inserted for vector construction (Fig. 5a). Subsequently, above two vectors were packaged into AAV-PHP.eb virus by Shanghai Obio corporation in order to efficiently infect with neuronal cells in the whole brain via crossing the BBB. The corresponding sequences used in the plasmid construction were listed as followed.

Human *MEF2C* L35P point mutation sequence:

forward: 5'-CGAGCGTGCTGTGTGACTGTGAGATTGC-3'.

reverse: 5'-GCTCATAAGCCTTCTTCATCAACCCAAATTTCCTC-3'.

Human *MEF2C* L35F point mutation sequence:

forward: 5’-TTCAGCGTGCTGTGTGACTGTGAGATT-3’.

reverse: 5’-GCTCATAAGCCTTCTTCATCAACCCAAATTTCCTC-3’.

Human *MEF2C* L35I point mutation sequence:

forward: 5’-ATAAGCGTGCTGTGTGACTGTGAGATT-3’.

reverse: 5’-GCTCATAAGCCTTCTTCATCAACCCAAATTTCCTC-3’.

Human *MEF2C* L35A point mutation sequence:

forward: 5’-GCCAGCGTGCTGTGTGACTGTGAGATT-3’.

reverse: 5’-GCTCATAAGCCTTCTTCATCAACCCAAATTTCCTC-3’.

Mouse *Mef2c* shRNA-1 sequence: 5’-GCCATCAGTGTCTGAGGATGT-3’.

Mouse *Mef2c* shRNA-2 sequence: 5’-GTCTGAGGATGTGGATCTGCT-3’.

The DsRed shRNA sequence was used as balnk control: 5’-AGTTCCAGTACGGCTCCAA-3’.

Mef2c sgC8 sequence: 5’-TATGAGCCGAGCGTGCTGTG-3’.

Mef2c sgC15 sequence: 5’-AAGGCTTATGAGCCGAGCG-3’.

**Animals.**

The C57BL/6 *Mus musculus* was used as the experimental model in this article. All procedures were approved by the Animal Care and Use Committee of the Center for Excellence in Brain Science & Intelligence Technology, Chinese Academy of Sciences, Shanghai, China. The use and care of animals were in accordance with the guidelines of this committee. According to animal welfare requirements, all experimental mice were bred in a pathogen free (PF) unit under constant temperature (approximately 22 ℃), humidity (approximately 55%RH), ventilation and automatic circadian rhythm on the condition of a 12h light/dark cycle (light from 7 a.m. to 7 p.m., dark from 7pm to 7 am) with food and water provided ad libitum. *Mef2c* L35P knock-in mice were designed and constructed by CRISPR/Cas9 mediated homologous recombination strategy on the genetic background of C57BL/6 mouse by Biocytogen Co.,Ltd. The sequences of small guide RNA (sgRNA) for *Mef2c* L35P knock-in mice and the targeting donor for *Mef2c* L35P knock-in mice were shown as followed. The genotype of *Mef2c* L35P knock-in mice, used in experiments was identified by sanger sequencing and tail genome PCR primers were shown as followed.

*Mef2c* L35P knock-in sgRNA sequence:

5'-ATATGTTCATTTACTGCGAC TGG-3'.

5'-ATGAAGGACTATATAGTCCA AGG-3'.

*Mef2c* L35P knock-in mice sequence:

forward: 5'-GAATGTGTTAGCACCCAAGACTCTG-3'.

reverse: 5'-GCATGTTGCAGCCATAGATGGGGTA-3'.

**Primary mouse cortical neuron culture.**

Mouse cortical neurons were extracted from 14.5 days embryos of either sex (Yu and Malenka, 2003). Cerebral cortices were dissected, digested with 20 U/ml papain (Worthington, LS003126) at 37 °C for 30 min, and cultured in 0.5 mL/well Neurobasal medium (Gibco, 21103-049) with 0.2% B27 (Gibco, 17504-044) and 2 mM Glutamax-I (Gibco, 35050-061) on Lab-Tek II Chamber Slide (Thermo Fisher Scientific, 154,941) at 100,000 cells/cm$^2$. Lipid transfection was used for investigating the variation of neuronal morphological development *in vitro*. After transfection, cultured neuronal cells were fed with new medium at interval of two days. For the neuronal morphology experiment, the cells were transfected by lipofectamine 2000 (Invitrogen, 11668-019) with 0.2 μg of each vector after 24 hours (hr) cultivation according to Lipofectamine® 2000 Reagent protocol. Neuronal cells were immunofluorescent stained at DIV4 for axonal morphology and at DIV10 for dendritic morphology, respectively. To be mentioned, Simple Neurite Tracer plugin in Image J software was utilized to quantify the total axonal length, axonal neurite length, dendritic branch numbers and dendritic branch length of cultured neurons. For the test of investigating molecular mechanism underlying the effect of L35P on MEF2C protein expression disruption, neuronal cells were transiently infected with lentivirus (dosage: $1.2\times10^{7}$ vg) 24 hr after planting and replaced medium 24hr after transfection. Cells were harvested at DIV5 for subsequent quantitative real-time PCR or western blotting assays. Besides, electroporation was conducted for identifying cytosine base editors (CBE) efficiency in primary cortical neurons *in vitro* and the neurons were electroporated with 3 μg of each plasmid mediated by specialized electroporation solution mixed of solution I, made of $C_{10}H_{14}N_5Na_2O_{13}P_3 \cdot H_2O$ (A2383, sigma) and $MgCl_2 \cdot 6H_2O$ (M2393, sigma), and solution II, made of $KH_2PO_4$ (p5655, sigma), $NaHCO_3$ (S5761, sigma) and glucose (G6152, sigma), at a proportion of 1:50.

**HEK293 cells culture and preparation of protein samples.**

HEK293 cells were cultured to approximately 90% proportion in culture dish (Corning) with the DMEM (Gibco, 11,965–092) medium and 10% FBS addition, and exogenously transfected with corresponding expression plasmids by using Lipofectamine 3000 (Invitrogen, L3000075) instructed by the manufacturer. After incubating for 72 hr, the cells were washed with 1×PBS, and subsequently harvested by gentle pipetting and lysed in loading buffer. HEK293 cells were heated at 100 °C for at least 30 mins to ensure the adequate denaturation of protein and then stored at -20°C. Similarly, the extraction of protein from cultured neuronal cells was manipulated by the same procedure as HEK293 cells.

**RNA extraction and reverse transcription.**

Total RNA was extracted from cultured mouse cortical neurons in Trizol reagent (Invitrogen, 15,596–018) and manipulated guided by TRIzol™ Reagent protocol. The reverse transcription was carried out with PrimeScript RT Master Mix (TaKaRa, RR036A) according to the corresponding protocol of manufacturer, and 500-1000ng total RNA was used as PCR template per reaction.

**Quantitative real-time PCR (qPCR).**

Quantitative real-time PCR was used for investigating the relative mRNA level of *Mef2c* normalized to GAPDH. For qPCR manipulation, the experiment was conducted with SYBR green premix (Toyobo, QPK-201) and data was analyzed through the StepOnePlus RealTime PCR System (Applied Biosystems). The qPCR program was set up three steps with melting as follows: 95 °C denaturation for 10 min, followed by amplification of 40 cycles of 95 °C for 10 s, 60 °C for 15 s, and 72 °C for 20 s. *Mef2c* relative mRNA expression level was calculated and standardized by means of the ΔCT method and *Gapdh* was regarded as internal control. The following primers were used in qPCR experiments:

mouse-*Mef2c*:

forward: 5’-GGATGGTAACTGGCATCTCAA-3’.

reverse: 5’-TGATCAGCAGGCAAAGATTG-3’.

mouse-*Gapdh*:

forward: 5’-ACGGCCGCATCTTCTTGTGCAGTG-3’.

reverse: 5’-GGCCTTGACTGTGCCGTTGAATTT-3’.

**Western blotting.**

The protein samples extracted from mouse tissues were lysed with RIPA buffer (containing 150 mM NaCl, 1% sodium deoxycholate, 0.1% SDS, 50 mM Tris-HCl pH 7.4, 1% TritonX-100 and protease inhibitor cocktail tablets (Roche, 04693159001)). The tissues were sufficiently digested on the rotated shaker at 4°C and the lysates were centrifuged at 12,000 rpm for 10 min at 4 °C. The supernatant of tissues lysates after centrifugation were collected as protein samples and then heated at 100 °C for at least 30 mins for adequate denaturation. Subsequently, the protein samples were conducted to run gel electrophoresis (10% SDS-PAGEs, Tris-Glycine system) at constant voltage of 80V for stack and 120V for separation, and then transferred to PVDF membranes (Millipore, pore size: 0.45 μm) at constant current of 200 mA. The transferred membranes carrying target protein were blocked in 5% BSA diluted with 1×TBS-Tween for 3 hr at room temperature and then incubated with primary antibodies overnight at 4 °C. After washing with 1×TBS-Tween, the membranes were incubated with secondary antibodies for 3 hr at room temperature. The protein bands were detected via chemiluminescence (ECL Western Blotting Substrate, Pierce, #32106).

**Fluorescence-activated cell sorting.**

Neuronal cells transfected with sgRNA-mCherry and CBE-EGFP plasmids were digested into a single-cell suspension and collected before acquisition on a flow cytometer at DIV4. Cells were sorted and counted on a BD FACScan flow cytometer (FACSAria II) (BD Biosciences). In addition, we used the 561- to 614-nm channel to detect mCherry signal and the 488- to 513-nm channel for EGFP. Data were acquired with BD CellQuest Pro software (BD Biosciences) and analyzed with FlowJo flow cytometry analysis software (Tree Star).

**Immunohistochemistry.**

For the immunofluorescent staining of cultured primary neurons, the medium of cells were disposed and then washed with 1×PBS. 4% paraformaldehyde (PFA) diluted with 1×PBS was used to fix the cells for 20 min at room temperature. After washed with 1×PBS, cells were incubated in the block buffer (5% BSA, 0.3% Triton X-100 diluted with 1×PBS) for 3 hr at room temperature. Subsequently, fixed neuronal cells were incubated with primary antibodies (5% BSA, 0.1% Triton X-100 diluted with 1×PBS) overnight at 4°C, followed by the incubation of secondary antibodies for 3 hr at room temperature. To acquire brain slices, animals were firstly perfused transcardially with 1×PBS then fixed with 4% PFA. After fixation in 4% PFA overnight, mouse brains were dehydrated in 15% sucrose solution (diluted with 1×PBS) followed by 30% sucrose solution (diluted with 1×PBS). Brains after dehydration were sliced up into 40 μm thickness by means of cryostats sectioning of Leica CM1950. After cryostats slicing, the sections were washed with 1×PBS and then incubated in the block buffer for 3 hr at room temperature. Then the sections were incubated with primary antibodies overnight at 4°C, followed by the incubation of secondary antibodies for 3 hr at room temperature. All images were respectively captured on Olympus VS120 high-throughput fluorescence microscope, Nikon NiE-A1 plus upright confocal fluorescence microscope and Nikon TiE-A1 plus inverted confocal fluorescence microscope.

**Antibodies.**

The primary antibodies used in this article and dilutions were listed as followed : anti-MEF2A+MEF2C (1:1000; No.ab64644; host species: rabbit; Abcam, Cambridge, UK); anti-GAPDH (glyceraldehyde-3-phosphate dehydrogenase) (1:10000; No.ab8245; host species: mouse; Abcam, Cambridge, UK); anti-GFP (green fluorescent protein) (1:1500; No.E022030; host species: rabbit; Earth Life Sciences Inc., California, USA); anti-MAP2 (1:1000; MAB3418, host species: mouse; Millipore, Massachusetts, USA); anti-PV (parvalbumin) (1:1000; No.MAB1572, host species: mouse; Millipore, Massachusetts, USA); anti-SST (Somatostatin) (1:500; No.SC74556, host species: mouse; Santa Cruz Biotechnology, Santa Cruz de la Sierra, Spain); anti-SpCas9 (1:500; NBP2-36440, host species: mouse; Novus Biologicals, Littleton, Colorado, USA); anti-RFP (1:800; 600-401-379, host species: rabbit; Rockland Immunochemicals, Philadelphia, Pennsylvania, USA); anti-DAPI (4',6-diamidino-2-phenylindole) (1:1000; No. 28718-90-3, Sigma-Aldrich, Missouri, USA).

**Stereotactic and tail intravenous injection of AAV virus.**

For sparse labelling to investigate neuronal spine development *in vivo*, the mixture of AAV-hSyn-Cre (0.05μl of $1{\sim}1.25\times10^{13}$ vg $ml^{-1}$, generated by Shanghai Oibo corporation) and AAV-hSyn-DIO-tdTomato (0.05μl of $2.9\times10^{12}$ vg $mL^{-1}$, generated by Shanghai Oibo corporation) was directly injected stereotactically into bilateral medial prefrontal cortex (mPFC) of continuously anaesthetized mice after glass pipette penetration. The stereotactic coordinates targeting to the mPFC region were shown as followed: anteroposterior (AP), +2.45 mm; mediolateral (ML), 0.3 mm; dorsoventral (DV), -1.0 mm. The mice were euthanized and perfused one month later after viral stereotactic injection.

For verifying the base repair efficiency of CRSIPR-Cas9-mediated APOBEC3A-CBE system *in vivo*, the mixture of AAV-hSyn-1-793-U6-sgC8 (0.25μl of $5.1\times10^{12}$ vg $ml^{-1}$, generated by Shanghai Oibo corporation) and AAV-hSyn-794-APOBEC3A$^{Y130F}$-1368 (0.25μl of $6\times10^{12}$ vg $mL^{-1}$, generated by Shanghai Oibo corporation) was directly injected stereotactically into unilateral hippocampus (Hip) of continuously anaesthetized *Mef2c* L35P$^{+/-}$ mice after glass pipette penetration, while AAV-hSyn-EGFP (0.5μl of $5\times10^{12}$ vg $ml^{-1}$, generated by Shanghai Oibo corporation) was injected stereotactically into contralateral Hip as negative control. The stereotactic coordinates targeting to the Hip region injected with APOBEC3A-CBE gene editing tool were shown as followed: anteroposterior (AP), -2.0; -2.0; -3.0; -3.0 mm; mediolateral (ML), 1.5; 1.5; 2.75; 3.4 mm; dorsoventral (DV), 1.25; 1.75; 3.0; 3.0 mm. The mice were euthanized and perfused one month later after viral stereotactic injection.

For investigating the effect of amelioration of aberrant phenotype in *Mef2c* L35P$^{+/-}$ mice after delivery of APOBEC3A-CBE system, AAV-PHP.eB-hSyn-EGFP, AAV-PHP.eB-hSyn-1-793-U6-sgC8 and AAV-PHP.eB-794-APOBEC3A$^{Y130F}$-1368 were packaged by Shanghai Obio corporation and titered through qPCR. Anaesthetized 1-month-old *Mef2c* L35P$^{+/-}$ mice were injected through tail vein with the mixture of AAV-PHP.eB-hSyn-1-793-U6-sgC8 (150μl of $4.9\times10^{12}$ vg $ml^{-1}$) and AAV-PHP.eB-793-APOBEC3A$^{Y130F}$-1368 (150μl of $4.4\times10^{12}$ vg $ml^{-1}$), while AAV-PHP.eB-hSyn-EGFP (300μl of $4.5\times10^{12}$ vg $ml^{-1}$) was injected into *Mef2c* WT and L35$^{+/-}$ mice with the same manipulation as negative control. We euthanized the mice at 10-12 weeks of age to examine ASD-related neurodevelopmental pathological phenotypes.

**Behavioral tests.**

All male mice performed in the behavioral tests were at the age of eight weeks and handled for 4 days prior to manipulation of behavioral tasks. The behavioral data were recorded and analyzed by Ethovision XT software (Noldus, Wageningen, Netherlands, RRID:SCR_000441) or the investigator blinded to the genotype of mice. As the pre-condition of the operation of behavioral tasks, all experimental mice were placed in the behavioral test room beforehand to habituate the external environment for at least 1 hr.

The behavioral apparatus was thoroughly cleaned and removed residual smell with deodorant before and after the beginning of each task trial. In addition, mice were allowed to freely move and explore apparatus during the behavioral task. Behavioral tests were all conducted in the period of 9 a.m. to 6 p.m., which was in the light phase of the light-dark cycle. Mild light intensity was adjusted to 50 lux applied in the open field, marble burying, self-grooming and scratching and home cage social reciprocal test, and moderate light intensity was adjusted to 80 lux used in the three chamber test and novel object recognition task, while maximum light intensity was adjusted to approximately 300 lux as aversive stimuli in the elevated plus maze and Barnes maze.

**Open field test.**

The open field test was conducted in a designed and professional behavioral apparatus made of 0.75 cm thick white plastic board at the specification of 40 (length) × 40 (width) × 40 cm (height). At the beginning of each trial, each mouse was singly placed into the center of the box and allowed to move freely to adequately explore the apparatus for 30 min. Video recordings were captured with a Da Hua high-definition video camera during the test. In terms of behavioral analysis and plotting, the total moved distance, the mean moved velocity, the track traces and the duration spent in the center zone (20 × 20 cm) of the box were analyzed by using Ethovision XT software (Noldus) and plotted by using Image J, and furthermore the data analysis was blinded to the genotype of each mouse.

**Self-grooming and scratching pattern.**

The video of self-grooming and scratching pattern was recorded in the engineered and specialized behavioral apparatus made of 0.75 cm thick white plastic board at the specification of 48 × 24 × 24 cm with one 0.75 cm thick dark lightproof plastic board inserted into the middle of the apparatus. One experimental mouse was placed into the left chamber and while another test mouse was placed into the right chamber of the apparatus and allowed to move freely within the whole chamber for 30 min. Video recordings were captured with a Da Hua high-definition video camera during the behavioral task and self-grooming and scratching time was quantified by using second chronograph, and besides the data analysis was blinded to the genotype.

**The social intruder test**

According to the previous reports, the individuals clinically diagnosed with autism spectrum disorder were considered to be commonly affected in social recognition, such as face recognition. The individuals with autism appeared impaired face hallmarks discrimination and delayed response to face presentation in the face characteristics recognition task, revealing that social recognition might be regarded as the important evidence of identity of ASD animal models [2,3].

To investigate social recognition of *Mef2c* L35P$^{+/-}$ mice, previous protocols that had been developed were modified and adopted in this article [4-6]. The task was performed in the animal home cage removed of feed trough, food, and water bottles. To be mentioned, each experimental mouse was feed separately for five consecutive days in a single cage for social isolation and acclimated in the experimental environment for 1 hr before the start of the task. The home cage social recognition task included social approach and familiarity session, and social novelty and recognition session. In the task, recognition between social partner mouse was conducted with 5 min inter-trial intervals. On the social approach and familiarity session, a stranger mouse (mouse 1; C57BL/6J adult male mice, 8 week) was placed into the cage for 3 min and allowed to freely explore and interact on the first trial. Subsequently, three consecutive trials with the interval of 5 min between each trial was repeated to allow familiarity to mouse 1. On the social novelty and recognition session, a novel stranger mouse (littermate to mouse 1) was introduced. For data analysis of cumulative sniffing time, interaction time was recorded when the test mouse performed preference for actively sniffing to stimuli mouse and when the experimental mouse physically contacted with stimuli mouse through nose actively. The quantification of recognition index was calculated as: (cumulative sniffing time of trial 5) − (cumulative sniffing time of trial 4). Video recordings were captured with a Da Hua high-definition video camera during the task and cumulative sniffing time was quantified by using second chronograph, and besides the data analysis was blinded to the genotype.

**The three-chamber test.**

The three-chambered apparatus was conducted in the behavioral apparatus made of 0.75 cm thick white plastic board at the specification of 60 × 40 × 30 cm. The dummy plates in the apparatus had 4 cm × 4 cm cut-out doors guaranteeing free movement between chambers. The two side chambers contained small iron cages for harboring social partner mice. One day before the test, mice were habituated to the apparatus with empty cages in both side chambers for 1 hr, while social partner mice (C57BL/6J adult male mice, 8 weeks) were placed in the iron cages for 1 hr in order to relieve tension and anxiety. To be mentioned, each trial of three chamber behavioral pattern contained three session, habituation, social approach and social novelty respectively. On the test day, each mouse was put into the center chamber of the apparatus and allowed to move freely to adequately explore all three chambers on the habituation session. On the social approach session, the same test mouse was placed into the center chamber and meanwhile a social partner mouse (mouse 1) was placed into the iron cage in either of the side chamber, and subsequently allowed to move freely to explore within the apparatus for 10 min. On the social novelty session, the same test mouse was returned to the center chamber again and meanwhile a novel stranger mouse (mouse 2) was placed into the other side chamber, and allowed to move freely for 10 min. The entire apparatus was wiped with deodorant between each trial to eliminate odor cues between animals. Video recordings were captured with a Da Hua high-definition video camera during the test and the time spent interacting with social partner mouse or empty cage (20 cm × 10 cm) and locomotion heat maps were analyzed utilizing Ethovision XT software (Noldus) and plotted by Image J, blinded to the genotype.

**Novel object recognition test.**

Novel object recognition task was conducted in the apparatus made of 0.75 cm thick white plastic board measuring 24 × 24 × 24 cm. One day before the test day, each mouse was placed into the behavioral apparatus separately for 1 hr to habituate the environment. On the habituation session of the task, each test mouse was placed into the center of the apparatus and allowed to move freely to adequately explore the entire apparatus for 10 min. On the training session one day after habituation session, test mouse was singly placed into the center of the apparatus and allowed to move freely to adequately explore for 10 min with two identical blue cylinders (object 1) on the two top corners of the apparatus. On the recognition session one day after training session, each test mouse was put into the center of the apparatus and allowed to move freely to adequately explore for 10 min with one blue cylinder (object 1) and one green cone (object 2) on the top corners of the apparatus. Video recordings were captured with a Da Hua high-definition video camera during the test and the time spent interacting with each object 1 or novel object (nose point range $\leq$ 3 cm) and locomotion heat maps were analyzed using Ethovision XT software (Noldus) and plotted by Image J, blinded to the genotype of each mouse.

**Elevated plus maze test.**

Elevated plus maze was conducted in the specialized behavioral apparatus, closed arm of which was made of 0.75 cm thick white plastic board measuring 30 × 5 × 15 cm, while open arm was designed at the specification of 30 × 5 × 1 cm and the height of entire apparatus was 50 cm. Each test mouse was placed into the central position of the plus maze and allowed to move freely to adequately explore the whole maze for 10 min. Video recordings were captured with a Da Hua high-definition video camera, and time spent in the open and closed arm and data analysis were performed through utilizing Ethovision XT software (Noldus) and plotted using Image J, and moreover the data analysis was blinded to the genotype of each mouse.

**Barnes maze spatial memory test.**

The Barnes maze was made of a rounded, thick, white plastic board with diameter of 122 cm. Forty holes with diameter of 5 cm were evenly spaced around the perimeter of the maze. The height of the maze above ground was 80 cm and designed to be rotated at the center of circle. The escape cage was made of a black lightproof plastic box and beneath the target hole for accessibility. Two bright supporting lamps placed on the opposite direction were adjusted to maximum light intensity as intensive aversive stimuli. The maze and escape cage were thoroughly cleaned and removed residual smell with deodorant between each training trial and the maze was rotated randomly after each trial to avoid any olfactory cue. To be mentioned, the Barnes maze test included training session for four consecutive days, followed by test session on the fifth day. Before the training session, mice were habituated to the environment for at least 1 hr. For each trial, each mouse was placed on the center of the maze, covered with a dark opaque plastic box for approximately 10 seconds (sec) and subsequently allowed to freely move to explore the entire maze. Each mouse was given 3 min at most to explore the maze to discover the location and direction of escape hole by itself, if not the mouse would be guided to the target hole and familiar with the direction of the target hole. Then, the mouse was allowed to stay in the box beneath the target hole for 2 min to acquisitive the rules. In addition, to be mentioned a dark opaque plastic lid would cover onto the box to avoid aversive hard light. Each mouse would go through four consecutive training trials in one day. Repeatedly, the mice were trained to establish memory of the target hole location (each mouse: 4 times per day, 16 times in total) and learn to orientate the target hole by themselves. On the test session, the escape box beneath the target hole was removed and each mouse was allowed to freely explore on the maze for 90 sec. Video recordings were captured with Da Hua Smart video recording software. The primary latency in the target or opposite zone was calculated within 90 sec after the beginning of test and locomotion track traces were analyzed via the Ethovision XT software (Noldus) and plotted by Image J, blinded to the genotype.

**List of Off-target primers**

OT1
Forward: CCTGAGACTTTCCATGGCTGACTG
Reverse: CTGCTGGTTGAGGACAGTACTTCTTC

OT2
Forward: TGTGCTGTAGGTGACTTTCACCAAG
Reverse: ACCTGAAGGATATCCGCGTTGG

OT3
Forward: CGGAGAGATACAGTGACACAGAGAG
Reverse: TGCCCTGTTCCCACTACGGAA

OT4
Forward: GGTTGGAAGGTGGAATGGCTAG
Reverse: GCTACCTCGTTAGTCCCTCATGGC

OT5
Forward: AGTGTGCCCAGGCAGATAGCA
Reverse: CCGCCTTACTGGTGGCCATAGA

OT6
Forward: CGTTCAGATCACAAGAAGGCAAGG
Reverse: CCCGTAAAGGGGTTGCAACAT

OT7
Forward: CCACACGATGCTCTCCTGCAG
Reverse: CTGGGTTCAGGAGAAGAGATGGGTTTA

OT8
Forward: AAAGGCATCCATCCCGTGTTT
Reverse: ACCCAGGGGGTTTACTTGGT

OT9
Forward: GCTTCGGGGTCTGTCTTAAGGG
Reverse: GGCTTCGTAGGCACTTTGCATT

OT10
Forward: GGTTTCTGCTCTTCGTCCTGTTCTT
Reverse: ATCCAGGCATTTGTCATTTGACTGT

OT11
Forward: TCCTTACAAGATGGGTGGGGCC
Reverse: AATGGCTTATAGATCGTTGGTGAGTCC

**Statistical analysis.**

All statistical data represented mean ± s.e.m., and *n* values indicate the number of individual experiments or mice. We used GraphPad Prism (v.6.01) and Origin (v.2019b) to analyze statistical variation among groups. The significance of differences among groups was assessed by the paired or unpaired two-tailed Student's *t*-test, one-way repeated-measures analysis of variance (ANOVA) and two-way ANOVA as indicated in the figure legends. To be mentioned, paired two-tailed t-test was performed to analyze the cumulative time interacting with mouse on the social approach and social novelty session in three chamber test within a single mouse. One-way, repeated-measures ANOVA was performed to analyze the difference between the means of more than two groups under the premise of single factor dependence and two-way ANOVA was performed to analyze under the premise of double factor dependence, and besides, the unpaired two-tailed t-test was performed in other instances. Statistically significance of intergroup differences was indicated at $*p < 0.05$, $**p < 0.01$ and $***p < 0.001$. Sample size were pre-assessed without any statistical testing methods. Mice were randomly allocated to different experimental groups. The immunohistochemical and behavioral tests were conducted blinded to the genotypes of the mice.

**Supplementary References**


1 Li, S. *et al.* Docking sites inside Cas9 for adenine base editing diversification and RNA off-target elimination. *Nat Commun* **11**, 5827, doi:10.1038/s41467-020-19730-9 (2020).

2 McPartland, J. C., Webb, S. J., Keehn, B. & Dawson, G. Patterns of visual attention to faces and objects in autism spectrum disorder. *J Autism Dev Disord* **41**, 148-157, doi:10.1007/s10803-010-1033-8 (2011).

3 Weigelt, S., Koldewyn, K. & Kanwisher, N. Face identity recognition in autism spectrum disorders: a review of behavioral studies. *Neurosci Biobehav Rev* **36**, 1060-1084, doi:10.1016/j.neubiorev.2011.12.008 (2012).

4 Hornberg, H. *et al.* Rescue of oxytocin response and social behaviour in a mouse model of autism. *Nature* **584**, 252-256, doi:10.1038/s41586-020-2563-7 (2020).

5 Jennifer N. Ferguson1, L. J. Y., Elizabeth F. Hearn1, Martin M. Matzuk2–4, Thomas R. Insel1 & James T. Winslow. Social amnesia in mice lacking the

oxytocin gene. *Nature Genetics* **25**, 284 (2000).

6 R. Dantzer, R.-M. B., G.F. Koob, and M. Le Moal. Modulation of social memory in male rats by neuropophseal peptides. *Psychopharmacology* 363 (1987).
